# Emerin self-assembly and nucleoskeletal coupling regulate nuclear envelope mechanics against stress

**DOI:** 10.1101/2021.02.12.429834

**Authors:** Anthony Fernandez, Markville Bautista, Liying Wu, Fabien Pinaud

## Abstract

Emerin is an integral nuclear envelope protein participating in the maintenance of nuclear shape. When mutated or absent, emerin causes X-linked Emery-Dreifuss muscular dystrophy (EDMD). To define how emerin takes parts in molecular scaffolding at the nuclear envelope and helps protect the nucleus against mechanical stress, we established its nanoscale organization using single molecule tracking and super-resolution microscopy. We show that emerin monomers form localized oligomeric nanoclusters stabilized by both lamin A/C and SUN1 at the LINC complex. Interactions of emerin with nuclear actin and BAF additionally modulate its membrane mobility and its ability to oligomerize. In nuclei subjected to mechanical challenges, the mechanotransducing functions of emerin are coupled to changes in its oligomeric state, and the incremental self-assembly of emerin determines nuclear shape adaptation against forces. We also show that the abnormal nuclear envelope deformations induced by EDMD emerin mutants stem from an improper formation of lamin A/C and LINC complex-stabilized emerin oligomers. These findings place emerin at the center of the molecular processes that regulate nuclear shape remodeling in response to mechanical challenges.

## Introduction

Emerin is an inner nuclear membrane (INM) protein that participates in the maintenance of the nuclear architecture by interacting with the lamina and elements of the linker of the nucleoskeleton and cytoskeleton (LINC) complex(Berk et al., 2013b; Kirby and Lammerding, 2018). Emerin also contributes to tethering chromatin at the nuclear envelope (NE) by binding the DNA-bridging barrier-to-autointegration factor (BAF) and regulating the activity of chromatin compaction modulators(Cui et al., 2015; Demmerle et al., 2012). Both the nucleoskeleton and the compaction state of chromatin contribute to the mechanical responses of nuclei and emerin is deemed a pivotal actor of mechanotransducing processes at the NE(Maurer and Lammerding, 2019). When mutated or absent, emerin causes X-linked Emery-Dreifuss muscular dystrophy (EDMD), a disease part of a larger of group of laminopathies associated with structural perturbations of the NE and its underlying lamina(Emery and Dreifuss, 1966). While emerin is ubiquitously expressed, mutations in the *EMD* gene primarily affect cells exposed to extensive mechanical stress, such as skeletal and cardiac muscle cells. Emerin-null muscle tissues display deformed and disorganized nuclei, impaired myogenesis and improper muscle fiber formation, which participate in the muscle wasting and cardiac disease phenotypes of EDMD(Fidzianska and Hausmanowa-Petrusewicz, 2003). In non-muscle cells, loss of emerin also induces altered NE elasticity and increased nuclear fragility(Rowat et al., 2006), impaired expression of mechanosensitive genes and enhanced apoptosis after mechanical strain(Lammerding et al., 2005). EDMD phenotypes might stem from both an altered structural integrity of the NE and modified gene expression profiles.

The structure of emerin and its binding to various partners have been extensively characterized(Berk et al., 2013b). Emerin is 254 amino acids (aa) long, with a N-terminal globular LAP2-emerin-MAN1 domain (LEM, aa:1-45), followed by a long intrinsically disordered region (IDR, aa:46-222) and a C-terminal transmembrane domain (aa:223-234) that allows anchoring to the NE. Prominent binding partners of emerin include lamin A/C and lamin B(Clements et al., 2000; Sakaki et al., 2001), actin(Fairley et al., 1999), BAF(Lee et al., 2001), nesprins and SUN proteins from the LINC complex(Haque et al., 2010; Mislow et al., 2002) and other factors(Berk et al., 2013b). While binding of emerin to BAF takes place via the LEM domain, interactions with other partners and emerin self-association are mediated by the IDR(Berk et al., 2014; Herrada et al., 2015). Emerin can indeed oligomerize via IDR segments that serve as LEM binding and self-assembly sites between emerin monomers, and, in vitro, emerin self-association into filamentous structures impacts binding to lamin A/C and BAF(Berk et al., 2014; Samson et al., 2017). Post-translational modifications of emerin also influence its oligomerization and its interactions with binding partners(Berk et al., 2013a; Roberts et al., 2006). In isolated nuclei, phosphorylation of emerin residues Y74 and Y95 are required for lamin A/C recruitment to the LINC complex and NE stiffening in response to mechanical loads(Guilluy et al., 2014). Additional post-translational modifications regulate the affinity of the LEM domain to BAF(Berk et al., 2013a) and binding to actin(Hirano et al., 2005). How emerin organizes at the NE and participates in protecting the nucleus against mechanical strains remains unclear.

EDMD often involves frame-shift deletions and nonsense mutations that effectively ablates emerin expression in cells. Yet, a few sets of point mutations and deletions, including Δ Q133H and P183H/T, also cause EDMD despite the localization of these mutated emerin at the NE and normal cellular expression levels(Fairley et al., 1999; Holt et al., 2001). These mutants display altered self-association and binding to nucleoskeletal proteins in vitro(Berk et al., 2013b; Herrada et al., 2015) and possible changes in their molecular organizations at the NE might explain why they induce EDMD.

Here, we used super-resolution imaging by direct stochastic optical reconstruction microscopy (dSTORM)(van de Linde et al., 2011) and single particle tracking by photoactivated localization microscopy (sptPALM)(Subach et al., 2010) to investigate the nanoscale organization of wild-type and mutant emerin in cells. By modulating the mechanical landscape of nuclei using cell micropatterning techniques(Fernandez et al., 2017), we also studied how emerin participates in NE mechanotransduction processes during stress. We show that emerin oligomerization and its differential interactions with key structural partners at the NE is central to nuclear shape adaptation against mechanical challenges.

## Results

### Emerin displays distinct diffusive behaviors at the nuclear envelope

To study the NE dynamics of wild-type and dystrophy causing mutations of emerin, PA-TagRFP-emerin fusions were expressed in emerin-null human dermal fibroblasts (HDF) from an EDMD patient (*EMD*^−/y^). PA-TagRFP-emerin expression led to its expected localization at the NE, as observed for endogenous emerin in HDF from a healthy individual (*EMD^+/y^*) (Fig. 1A). Upon re-expression of wild-type emerin fusions in *EMD*^−/y^ HDF, nuclear deformations against mechanical stress recovered to levels similar to wild-type *EMD*^+/y^ HDF(Bautista et al., 2019). For sptPALM, individual PA-TagRFP-emerin were imaged using highly inclined and laminated optical sheet excitation at the ventral NE(Tokunaga et al., 2008). Diffusion tracks were generated from localized emerin appearances (Fig. 1B; Movie S1) and analyses were performed using the probability distribution of square displacements (PDSD)(Schutz et al., 1997) to separate unique diffusive behaviors (Fig. 1C).

**Fig. 1.**
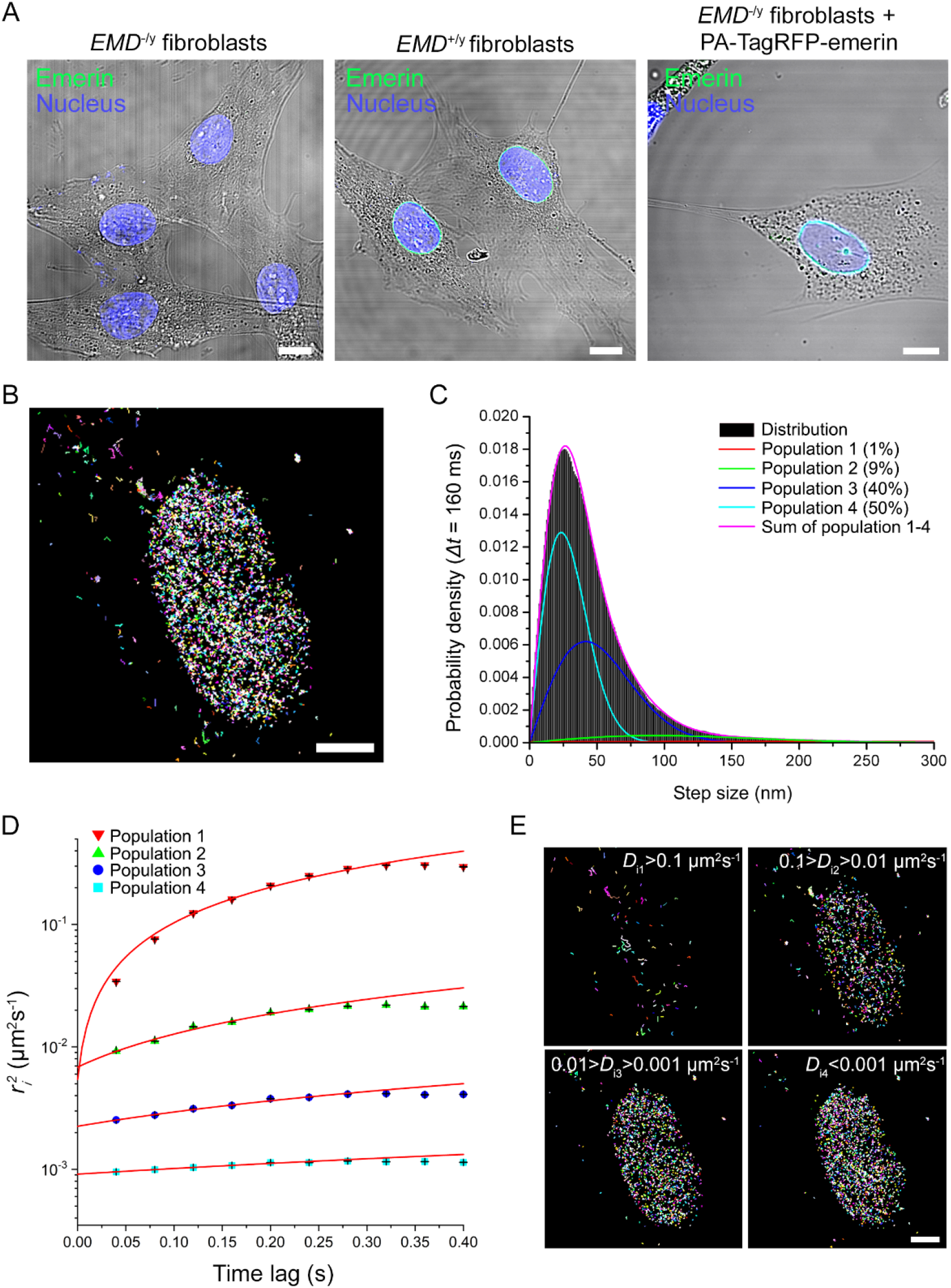
Emerin displays distinct diffusive behaviors at the nuclear envelope. (A) Confocal imaging of emerin in *EMD*^−/y^ HDF, *EMD*^+/y^ HDF and *EMD*^−/y^ HDF expressing wild-type PA-TagRFP-emerin. Scales: 10 μm. (B) Trajectory map of individual PA-TagRFP-emerin at the NE. Scale: 5 μm. (C) Probability distribution of square displacements for wild-type emerin at time lag Δ*t* = 160 ms. (D) Square displacement curves (r^2^*_i_*) of each emerin diffusive behavior fitted over he first four values with a Brownian diffusion model (red line). (E) Emerin trajectory maps as a function of individual diffusion coefficient (*D*_i_). Scale: 5 µm.

We observed that wild-type emerin exhibits four distinct diffusive behaviors with diffusion coefficients D_1_: 2.21×10^−1^ ± 4.9×10^−2^ µm^2^s^−1^, a value similar to that reported for ER membrane diffusion(Ostlund et al., 1999), D_2_: 1.48×10^−2^ ± 1.5×10^−3^ µm^2^s^−1^, D_3_: 1.73×10^−3^ ± 1.1×10^−4^ µm^2^s^−1^, and D_4_: 2.6×10^−4^ ± 1×10^−5^ µm^2^s^−1^ (Fig. 1D; Table S1). The ER population D_1_ makes up 1% of the detected behaviors, while population D_2_ represents 9%, a small emerin fraction comparable to that expected at the outer nuclear membrane (ONM)(Salpingidou et al., 2007). The two slowest populations D_3_ and D_4_ collectively represent 90% of emerin diffusive behaviors. Based on their individual diffusion coefficient, we grouped emerin trajectories into four diffusion ranges and plotted them as maps. In trajectory maps, the fastest emerin population D_1_ primarily distributes in the vicinity of the NE, consistent with diffusion in the ER membrane, while the three slower populations D_2-4_ are enriched at the NE (Fig. 1E). Compared to measurements by fluorescence recovery after photobleaching, where only two diffusive behaviors of emerin were detected (Fig. S1), sptPALM reveals that emerin has four distinct types of diffusions.

### Emerin organizes as slowly diffusing monomers or oligomers at the inner nuclear membrane

While emerin primarily localizes at the INM, it is often detected at the ONM, where it does not interact with the nuclear lamina. To assess if diffusing population D_2_ corresponds to ONM- associated emerin, sptPALM was done after lamin A/C depletion by siRNA (Fig. S2). Downregulating the expression of lamin A/C did not affect emerin populations D_1_ and D_2_ (*p* not significant, Fig. 2A) but the mobility of populations D_3_ (40%) and D_4_ (50%) increased (*p*<0.01, Fig. 2A; Table S1), consistent with their localization at the INM and with population D_2_ diffusing at the ONM. Commensurate with the majority of emerin being distributed as two distinct INM pools, nuclear actin depletion by downregulating the expression of the actin nuclear import factor importin-9(Dopie et al., 2012) (IPO9, Fig. S2-S3) also resulted in the increased mobility of populations D_3_ and D_4_, although faster emerin diffusion at the ONM was also observed, for equivocal reasons (Fig. 2A; Table S1). The 9% fraction of emerin diffusing at the ONM agreed well with quantitative immunostaining measurements indicating that 13±6% of the emerin pool localizes at the ONM in *EMD^+/y^* HDF (Fig. S4).

**Fig. 2.**
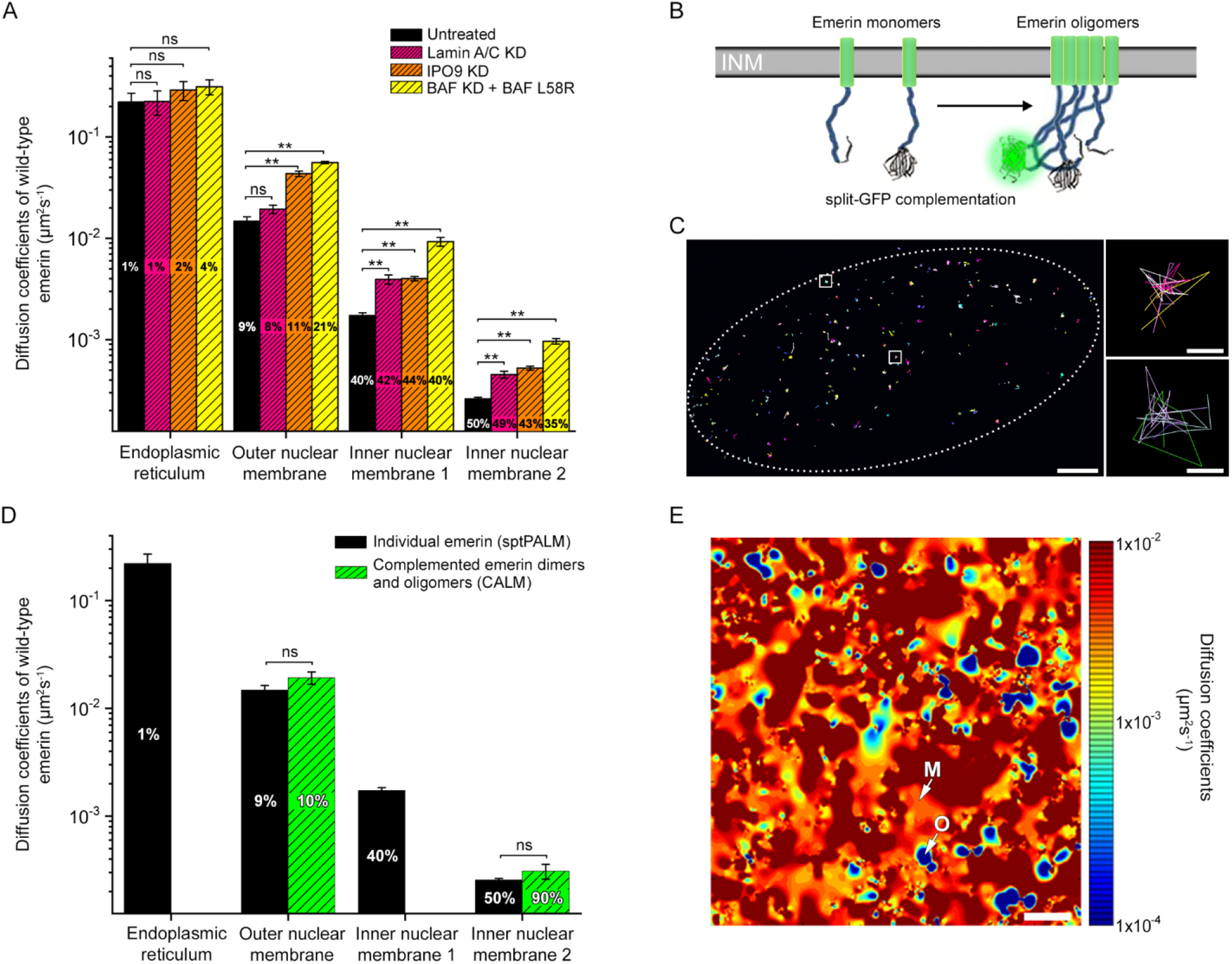
Emerin diffuses as monomers and oligomers interacting with lamin A/C, nuclear actin and BAF. (A) Diffusion coefficients (±s.e.m.) and percentages of wild-type emerin after lamin A/C depletion, nuclear actin depletion (IPO9 KD) or replacement of endogenous BAF with BAF^L58R^. (B) Schematic of emerin fusion to complementary split-GFP fragments. (C) Trajectory map of complemented emerin-GFP-emerin at the NE. Squares: oligomeric nanodomains with overlapping trajectories. Scales: 2 µm (left) and 50 nm (right). (D) Diffusion coefficients (±s.e.m.) and percentages for wild-type emerin assessed by sptPALM and complemented emerin-GFP-emerin species assessed by CALM. (E) Diffusion map of wild-type PA-TagRFP-emerin, showing slow mobility oligomeric domains (blue, O) surrounded by areas where emerin monomers diffuse faster (red, M). Scale: 500 nm. For (A, D): t-test, ns: non-significant, **: *p*<0.01.

Emerin can self-assemble in vitro, which could induce its slow INM mobility if emerin oligomers are tethered by the nucleoskeleton. To determine if diffusing populations D_3_ or D_4_ correspond to emerin monomers or oligomers, we co-expressed wild-type emerin fused to self-complementary split-GFP fragments (Fig. 2B), induced the formation of complemented emerin-GFP-emerin species and tracked their mobility by complementation activated light microscopy (CALM)(Pinaud and Dahan, 2011). We found that emerin-GFP-emerin species localize almost exclusively at the NE (Fig. 2C; Movie S2), with 90% diffusing like population D_4_ (*p* not significant, Fig. 2D) and 10% diffusing like the ONM population D_2_ (*p* not significant, Fig. 2D). No fluorescent species with mobility matching those of the ER population D_1_ or the INM population D_3_ were detected. Emerin-GFP-emerin trajectories often spatially overlap, suggesting that emerin forms multimers rather than strictly dimers at the INM (Fig. 2C). This indicates that population D_3_ detected by sptPALM but not by CALM represents emerin monomers, while the slowest population D_4_ represents INM emerin oligomers, as subsequently confirmed by super-resolution imaging. The detection of a few emerin-GFP-emerin species at the ONM is likely due to split-GFP-induced emerin dimers that cannot translocate through peripheral nuclear pore channels and do not access the INM.

### BAF binding modulates the mobility of both emerin monomers and oligomers distributed across the inner nuclear membrane

BAF binds the LEM domain of emerin with high affinity and could participate in the slow INM diffusion of emerin via its additional interactions with lamin A/C(Samson et al., 2018) and chromatin(Burger et al., 2020). To study how binding of emerin to BAF influences its mobility, endogenous BAF was depleted by siRNA (Fig. S2) and replaced by the expression of BAF^L58R^, a mutant that does not bind LEM domains but binds lamin A/C and chromatin(Samwer et al., 2017). When unable to bind BAF^L58R^, emerin monomers and oligomers diffuse significantly faster at the INM, with lateral mobilities higher than for lamin A/C or IPO9 knockdown (*p*<0.01, Fig. 2A; Table S1). While emerin diffusion at the ER is undisturbed, its ONM mobility increases (*p*<0.01, Fig. 2A), suggesting that cytoplasmic BAF also interacts with emerin at the ONM. BAF being highly mobile in the nucleus(Shimi et al., 2004), the comparatively slow mobility of BAF-bound emerin likely stems from the formation of ternary complexes with lamin A/C or chromatin. As knockdown of endogenous BAF and expression of BAF^L58R^ did not alter the expression of lamin A/C nor its nuclear localization (Fig. S2), the stronger influence of BAF^L58R^ on emerin dynamics as compared to lamin A/C depletion indicates that BAF not only modulates the mobility of emerin via ternary binding to lamin A/C, but also via additional interactions with other nuclear components, potentially chromatin.

The global influence of BAF and the nucleoskeleton on the mobility of emerin was also captured in diffusion maps built by spatial rendering of diffusion coefficients at the position of each tracked PA-tagRFP-emerin (Fig. 2E). In those map, slow diffusion domains attributed to emerin oligomers are distributed throughout the NE and are surrounded by areas where the mobility of emerin is faster, as expected for monomers (Fig. 2E). This indicates a membrane-wide, yet locally structured distribution of emerin, with monomers surrounding oligomer domains, consistent with emerin monomer/oligomer exchanges in distinct INM areas. Together, these results show that emerin diffuses rapidly at the ER membrane on its way to the juxtaposed ONM, where its diffusion is slower, in part by interaction with cytoplasmic BAF. Once at the INM, the mobility of emerin is further reduced, with monomers and oligomers interacting with nuclear BAF and the nucleoskeleton, consistent with a combined BAF-induced and lamina-induced retention of emerin at the INM.

### Emerin forms discrete oligomeric nanodomains surrounded by monomers

We then probed the organization of emerin in greater detail using super-resolution microscopy. Wild-type emerin fused to a SNAP-tag was expressed in *EMD*^−/y^ HDF and imaged at the ventral NE by dSTORM with benzylguanine-AlexaFluor647 (Fig. 3A). Emerin neighborhood density functions (NDF) built from localized emerin positions were compared to MonteCarlo simulations of completely random distributions and fitted to measure molecular densities and length scales of significant clustering. We found that wild-type emerin is not distributed randomly at the NE, as it displays density distributions significantly higher than expected for a completely random organization (Fig. 3B). NDF were best fitted with a two-exponential decay model, indicating that emerin organizes into two distinct clustering states across the NE. The first clusters have typical sizes of 22±11 nm and molecular densities of emerin 8.2±0.2 fold higher than expected for a random distribution (Fig. 3C; Table S2). The second clusters are significantly larger with typical sizes of 236±30 nm and emerin densities slightly above the expected value of 1 for complete randomness (1.3±0.1 fold, Fig. 3C). In local cluster maps, emerin forms small high-density clusters bordered by larger NE areas with emerin densities close to random (Fig. 3D). Those distributions are similar to sptPALM diffusion maps and corroborate that emerin organizes into discrete oligomers surrounded by dispersed emerin monomers across the INM. A similar organization of emerin was observed at the dorsal NE (Fig. S5). While it is intriguing that emerin monomers appear slightly clustered, their completely random distribution is not expected because they do not populate membrane-less areas of the NE, such as nuclear pores.

**Fig. 3.**
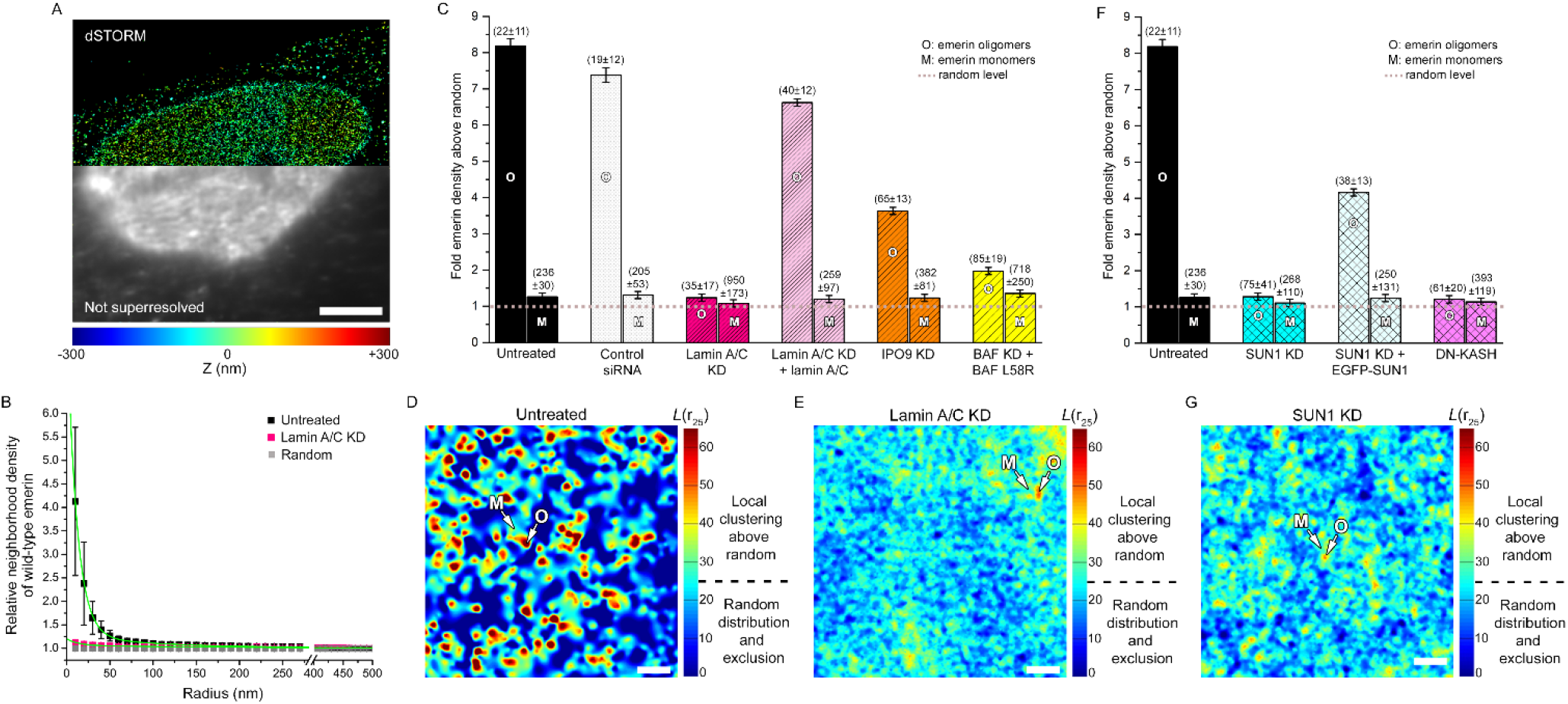
Emerin forms oligomeric nanodomains stabilized by lamin A/C and SUN1 at the LINC complex. (A) Super-resolved (top) and diffraction-limited imaging (bottom) of wild-type SNAP-emerin. Scale: 5 µm. (B) Neighborhood densities (±s.d.) at various length scales for wild-type emerin before and after lamin A/C knockdown, compared to complete spatial randomness. Fit shown in green. (C) Molecular densities above random (±s.e.m.) of wild-type emerin oligomers (O) and monomers (M) in untreated cells, after control siRNA, lamin A/C knockdown, lamin A/C knockdown and exogenous expression of lamin A/C, IPO9 knockdown to deplete nuclear actin or replacement of endogenous BAF by BAF^L58R^. Values in parenthesis represent the size (±s.e.m.) of each domain in nanometers. (D) Local cluster map of wild-type emerin at the NE of an untreated HDF. (E) Local cluster map of wild-type emerin after lamin A/C knockdown. (F) Molecular densities above random (±s.e.m.) for wild-type emerin in untreated cells, after SUN1 knockdown, after SUN1 knockdown and exogenous expression of EGFP-SUN1 or after exogenous expression of mCherry-DN-KASH. (G) Local cluster map of wild-type emerin after SUN1 knockdown. For (D, E, G), M: monomer areas, O: oligomer nanodomains. Scales: 250 nm.

### Emerin oligomers are stabilized by lamin A/C and SUN1 at LINC complexes and are modulated by nuclear actin and BAF

We then determined how lamin A/C, nuclear actin, BAF and SUN1 from the LINC complex participate in the spatial organization of emerin at the INM. Wild-type emerin was imaged by dSTORM after siRNA knockdown of lamin A/C, IPO9, SUN1, or replacement of endogenous BAF with BAF^L58R^. After lamin A/C depletion, the molecular density of emerin is reduced (Fig. 3B) and emerin distributes into 35±17 nm nanodomains with densities close to random (1.2±0.1 fold, Fig. 3C; Table S2) and into large 950±173 nm monomer domains with densities just above random (1.1±0.1 fold, Fig. 3C). Compared to controls, emerin monomers are dispersed over larger areas and oligomers nearly vanish, as seen in emerin cluster maps of lamin A/C-depleted nuclei (Fig. 3E). The spatial distribution of emerin and oligomerization returned to near normal levels upon exogenous expression of a siRNA-resistant lamin A/C (Fig. 3C and S6), but overexpression induced a loss of emerin oligomers (Fig. S6), indicating that balanced levels of lamin A/C are essential to promote emerin self-assemblies. Although lamin A/C depletion indirectly led to decreased nuclear actin levels (Fig. S2), emerin oligomers were not as severely disrupted when nuclear actin was specifically depleted, as shown next. These results indicate that lamin A/C plays a central role for the formation and the stabilization of emerin oligomers and additionally influences the spatial distribution of emerin monomers at the INM.

When nuclear actin is depleted after IPO9 knockdown, emerin forms oligomers having reduced molecular densities of 3.6±0.1 fold above random and sizes of 65±13 nm (Fig. 3C; Table S2). Emerin monomers are dispersed over 382±81 nm domains, larger than in untreated cells (236±30 nm, Fig. 3C), but not as large as for lamin A/C depletion (950±173 nm, Fig. 3C). Thus, despite reduced nuclear actin levels, emerin still oligomerizes but redistributes over broader INM domains. Conversely, nuclear accumulation of actin by siRNA of exportin 6 (XPO6, Fig. S3) did not affect the organization of emerin monomers, but oligomer densities decreased to 1.5±0.1 fold above random in 49±16 nm nanodomains (Fig. S7), suggesting that excess nuclear actin impedes the formation of dense emerin oligomers. Thus, together with lamin A/C, nuclear actin contributes to the spatial organization of emerin at the INM and participates in the maintenance of emerin oligomers.

When endogenous BAF is replaced by BAF^L58R^, emerin oligomers are still formed but with reduced molecular densities of 2.0±0.1 fold above random and larger sizes of 85±19 nm compared to controls (Fig. 3C; Table S2). The inability to bind BAF also induces a redistribution of emerin monomers over large INM domains with typical sizes of 718±250 nm (Fig. 3C). Consistent with our diffusion results, emerin binding to BAF modulates the organization of both oligomeric and monomeric emerin and strongly influences the spatial distribution of emerin monomers at the INM.

When SUN1 is depleted to destabilize LINC complexes (Fig. S2), the formation of emerin oligomers is reduced to levels equivalent to lamin A/C depletion. Emerin distributes over 75±41 nm nanodomains with molecular densities decreased to 1.2±0.1 fold above random (Fig. 3F; Table S2), and over NE areas with sizes of 268±110 nm, similar to the spatial distribution of emerin monomers in untreated cells (Fig. 3F). This indicates that reduced SUN1 expression levels primarily influence the formation of emerin oligomers at the INM. In cluster maps, depletion of SUN1 effectively results in a largely random distribution of emerin and, like for lamin A/C depletion, few emerin clusters are visible (Figure 3G). This loss of emerin oligomers was partially rescued upon expression of an exogenous and siRNA-resistant EGFP-SUN1 (Fig. 3F and Fig. S6), but overexpression induced a mis-localization of both EGFP-SUN1 and emerin from the NE (Fig.S6) and a concomitant reduction in emerin oligomerization (Fig. S6), indicating that, as for lamin A/C, a balanced cellular expression of SUN1 is required for the self-assembly of emerin at the INM. While SUN1 interacts with lamin A(Crisp et al., 2006; Haque et al., 2006), the expression and NE localization of lamin A/C was not disrupted by SUN1 knockdown and, reciprocally, lamin A/C depletion did not modify the expression of SUN1 nor its distribution at the NE (Fig. S2). Thus, together with lamin A/C, a regulated expression of SUN1 is required for the formation of emerin oligomers and their localized distribution at the INM. SUN1, itself, was shown to self-assemble into nearly immobile platforms that could serve as sites for macromolecular assemblies at LINC complexes(Hennen et al., 2018; Lu et al., 2008). To assess if emerin locally oligomerizes at LINC complexes, we expressed a dominant negative KASH domain of nesprin-1α (mCherry-DN-KASH)(Lombardi et al., 2011), which disrupts LINC complexes by binding to SUN proteins and impeding their interaction with endogenous KASH-domain proteins at the NE. In cells expressing mCherry-DN-KASH, emerin oligomers were disrupted at levels equivalent to those of SUN1 siRNA (Fig. 3C; Table S2), indicating that the oligomerization of emerin requires a functional coupling of the nucleoskeleton and the cytoskeleton via SUN1/nesprin interactions at LINC complexes.

Together, these results show that emerin monomers are distributed across the INM where their distribution is modulated by direct binding to BAF and additional spatial constrains imposed by lamin A/C and nuclear actin. Emerin also self-assembles into discrete and small oligomeric nanodomains that are structurally co-stabilized at the LINC complex by lamin A/C and SUN1 and are maintained, to a lesser extent, by BAF and nuclear actin.

### Increased diffusion and oligomerization of emerin upon nuclear adaptation to mechanical stress

We then studied how the mobility of emerin and its organization change in nuclei subjected to increasing mechanical stress. SptPALM and super-resolution imaging of wild-type emerin were done in *EMD*^−/y^ HDF grown on increasingly narrow rectangular micropatterns with widths of 15-5 µm to impose steady-state mechanical stress to the nucleus (Fig. 4A-B) and increase mechanical loads at LINC complexes(Arsenovic et al., 2016). First, we verified that changes in nuclear shape index (NSI) on these micropatterns reflect the mechanical adaptation of nuclei against forces by depleting lamin A/C or nuclear actin, two key nucleoskeletal proteins involved in maintaining nuclear shape. In non-patterned cells, nuclei are less circular after lamin A/C or nuclear actin depletion compared to controls (*p*<0.01, Fig. 4C). These effects are amplified on micropatterns where nuclei become increasingly deformed as patterns get narrower (*p*<0.01, Fig. 4C), indicating that cell micropatterning elicits increased nuclear stress and effectively induces mechanical responses that implicate the nucleoskeleton.

**Fig. 4.**
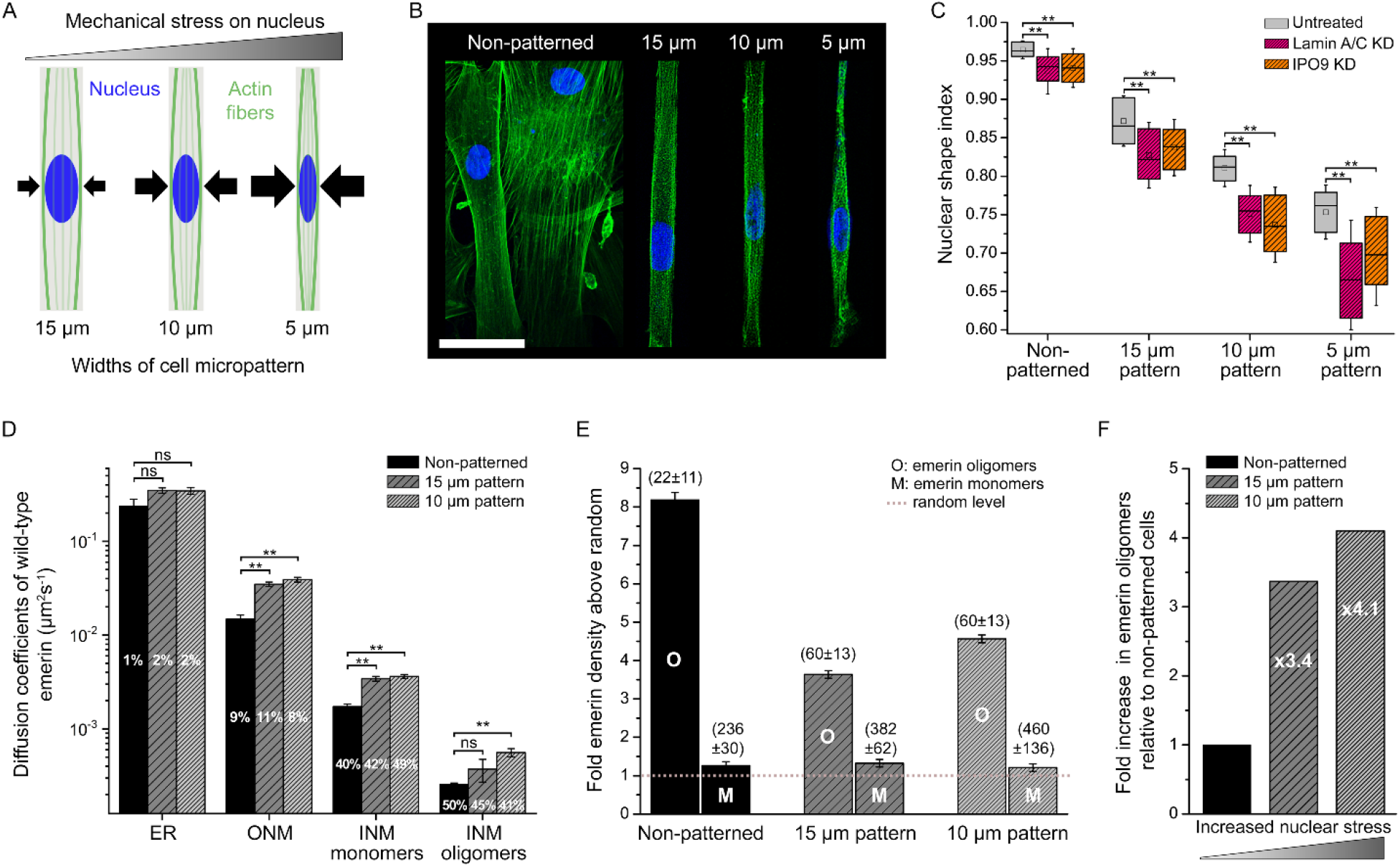
Mechanical stress increases emerin mobility and the formation of emerin oligomers. (A) Schematic of increasing nuclear mechanical stress by cell micropatterning. Arrows represent force. (B) Confocal imaging of actin (green) and the nucleus (blue) in micropatterned *EMD^+/y^* HDF. Scale: 50 µm. (C) Nuclear shape index as a function of micropattern width in *EMD^+/y^* HDF depleted for lamin A/C or nuclear actin. Wilcoxon test, **: p<0.01. (D) Diffusion coefficients (±s.e.m.) and percentages of wild-type emerin in non-patterned cells or after nuclear deformation on 15 and 10 µm micropatterns. t-test, ns: non-significant, **: *p*<0.01. (E) Molecular densities above random (±s.e.m.) for wild-type emerin oligomers (O) and monomers (M) in non-patterned *EMD^−/y^* HDF and after nuclear deformation on 15 and 10 µm micropatterns. Values in parenthesis represent the size (±s.e.m.) of each domain in nanometers. (F) Increase in emerin oligomers as a function of nuclear stress.

When we tracked wild-type PA-tag-RFP-emerin on deformed nuclei in 15 µm and 10 µm patterns (Movie S3), we found that its ER mobility is unchanged compared to non-patterned cells (*p* not significant, Fig. 4D; Table S1), while its diffusion at the ONM is faster (*p*<0.01, Fig. 4D). At the INM, emerin monomers also diffuse significantly faster (*p*<0.01, Fig. 4D) and the mobility of oligomers increases, albeit short of statistical significance for 15 µm patterns (*p* not significant and *p*<0.01, Fig. 4D). Interestingly, there was no significant difference in the mobility of all four ER, ONM and INM emerin populations in both types of micropatterns compared to nuclear actin depletion (*p* not significant, Fig. S8), suggesting that nuclear shape adaptation to mechanical cues entails modified interactions of emerin with nuclear actin, notably for monomers.

Spatial distribution analyses further indicate that, as the NSI decreases, emerin monomers disperse over increasingly large INM domains with sizes of 382±62 nm and 460±136 nm, compared to 236±30 nm for non-stressed nuclei (Fig. 4E). Concurrently, oligomer densities drop from 8.2±0.2 fold to 3.6±0.1 fold above random in 15 µm patterns, before increasing to 4.6±0.1 fold above random in 10 µm patterns (Fig. 4E). Remarkably, oligomeric nanodomains become larger during nuclear stress and their size expands from 22±11 nm to 60±13 nm in both micropatterns (Fig. 4E). When considering this wider spatial distribution, the relative oligomerization of emerin compared to non-deformed nuclei increases by 3.4 fold and by 4.1 fold as the nucleus adapts to incremental mechanical stress (Fig. 4F). This shows that nuclear shape adaptation to forces is associated with a gradual change in the oligomerization potential of emerin at the INM. This increased self-assembly of emerin over larger nanodomains is accompanied by a faster lateral mobility of emerin and is triggered by nucleoskeletal re-arrangements. Indeed, the stress-induced spatial reorganizations of emerin in micropatterns, including oligomer densities and emerin domain sizes, are analogous to those seen when nuclear actin is depleted in non-patterned cells (Fig. S8; Table S2). Consistent with the contribution of nuclear actin to changes in nuclear volume(Baarlink et al., 2017), these observations suggest that nuclear shape deformation involves a disengagement of nuclear actin from the nucleoskeleton that results in faster emerin diffusion and increased oligomerization. Increased lateral mobilities of emerin at the INM could indeed facilitate molecular collisions between monomers and the formation of oligomers stabilized by lamin A/C and SUN1 at LINC complexes. These results indicate that the mechanotransducing functions of emerin are coupled to changes in its oligomeric state along the INM and are modulated by emerin interactions with nucleoskeletal partners, including nuclear actin, to ensure appropriate nuclear deformation and response to mechanical challenges.

### Emerin mutants induce abnormal nuclear deformation against mechanical stress

To further establish the significance of emerin oligomerization for nuclear adaptation to mechanical stress, we then compared the organization of wild-type emerin with that of mutated forms of emerin known to induce EDMD. We studied how mutation Q133H, deletion Δ and mutation P183H (Fig. 5A) affect the diffusion and the nanoscale distribution of emerin at the NE. First, we verified that these mutated emerin effectively induce defective mechanical responses of nuclei when expressed in *EMD^−/y^* HDF by comparing changes in NSI after random cell plating or plating on increasingly narrow micropatterns (Fig. 5B-C). In randomly plated cells, the expression of mutated emerin results in slightly less circular nuclei compared to wild-type emerin (Fig. 5B). With increasing mechanical stress, cells expressing emerin mutants display significantly higher NSI than wild-type (Fig. 5B), indicative of a failure to correctly modify the shape of the nucleus in response to forces. These deficient changes in nuclear shape are accompanied by a mispositioning of the nucleus relative to the cell major axis, nucleus crumpling, abnormal organization of the actin cytoskeleton and failure of cells to properly fit within micropatterns, specifically in cell areas adjacent to the misshaped nucleus (Fig. 5C). It indicates that expression of mutated emerin in *EMD^−/y^* HDF effectively impedes nuclear adaptation to mechanical stress.

**Fig. 5.**
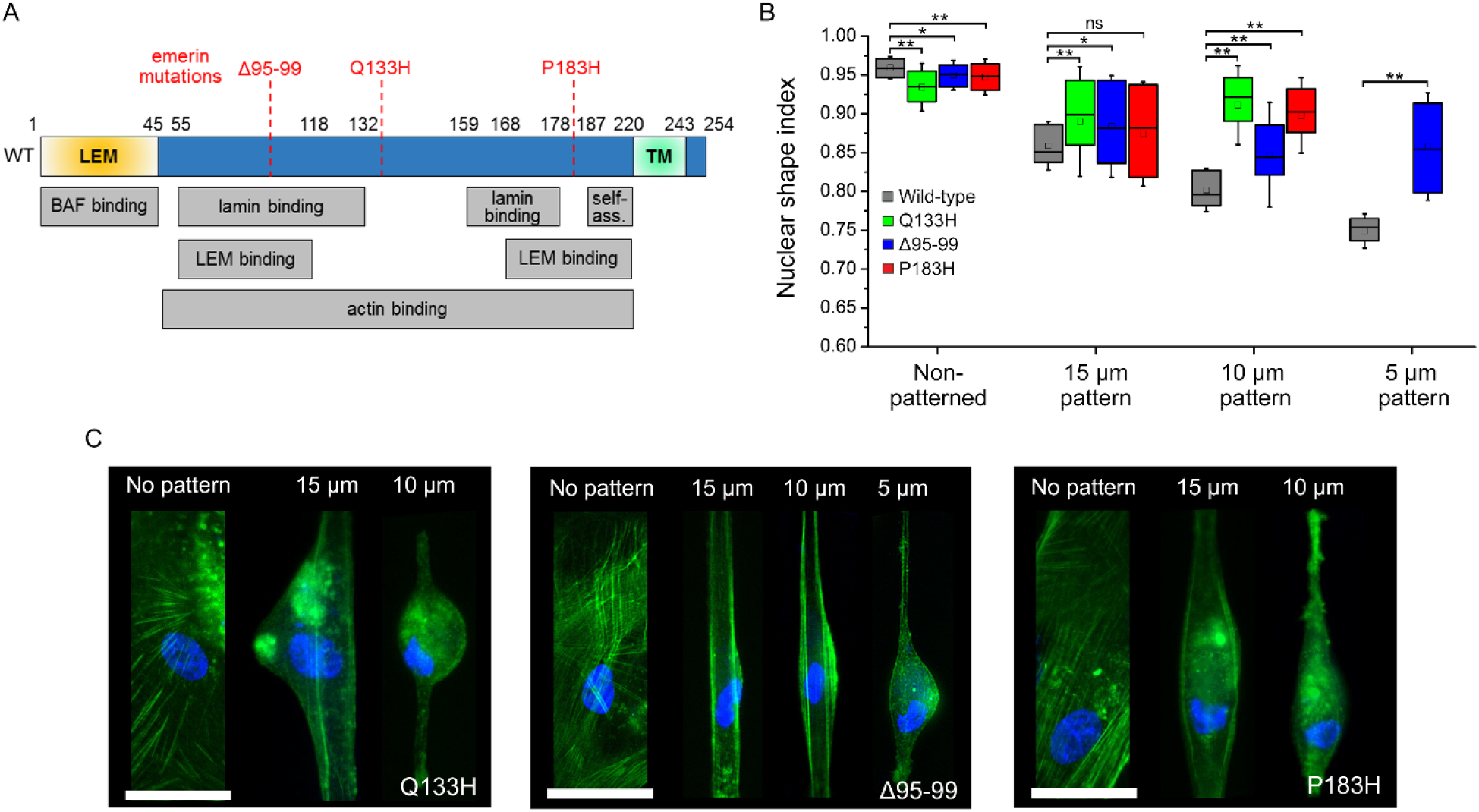
Emerin mutations induce defective nuclear shape adaptation against mechanical stress. (A) Diagram of emerin with binding and self-association domains (self-ass.) and position of Δ95-99, Q133H and P183H mutations. (B) Nuclear shape index as a function of micropattern width for *EMD^−/y^* HDF expressing wild-type, Q133H, Δ95-99 or P183H emerin. Wilcoxon test, ns: non-significant, *: *p*<0.05, **: *p*<0.01. (C) Fluorescence imaging of actin (green) and the nucleus (blue) in micropatterned *EMD^−/y^* HDF expressing Q133H, Δ95-99 or P183H emerin Scales: 50 μm.

### Emerin mutants display defective oligomerization at the inner nuclear membrane

#### Q133H mutation

We then characterized the NE dynamics of each mutated emerin and their respective nanoscale organization, starting with Q133H emerin. We found that the lateral mobility of Q133H PA-TagRFP-emerin is similar to that of wild-type emerin at the ER and the ONM (*p* not significant, Fig. 6A; Table S1), but that both Q133H monomers and oligomers diffuse significantly faster at the INM (*p*<0.01 and *p*<0.05, Fig. 6A). Previous in vitro studies indicated that the Q133H mutation disrupts emerin binding to actin(Holaska et al., 2004) but does not impede interactions with lamin A/C(Holt et al., 2001), SUN1(Haque et al., 2010) or BAF(Bengtsson and Wilson, 2004). The increased INM diffusion of Q133H, therefore suggests that it does not bind nuclear actin. Interestingly, this faster INM diffusion of Q133H resembles the increased mobility of wild-type emerin when nuclear actin is depleted (Fig. 2A), which further underlines that nucleoskeletal actin modulates the diffusion of emerin monomers and oligomers. This is consistent with previous observations that emerin expression influences the mobility of nuclear actin(Ho et al., 2013) and indicative of a reciprocal effect of emerin/nuclear actin interactions on their respective mobility. The diffusion of Q133H at the INM is also similar to that of wild-type emerin under mechanical stress, in both 15 µm and 10 µm wide micropatterns (*p* not significant, Fig. S8), which supports a mechanism where nuclear deformations involve a dissociation of emerin from nuclear actin that leads to faster emerin diffusion at the INM.

**Fig. 6.**
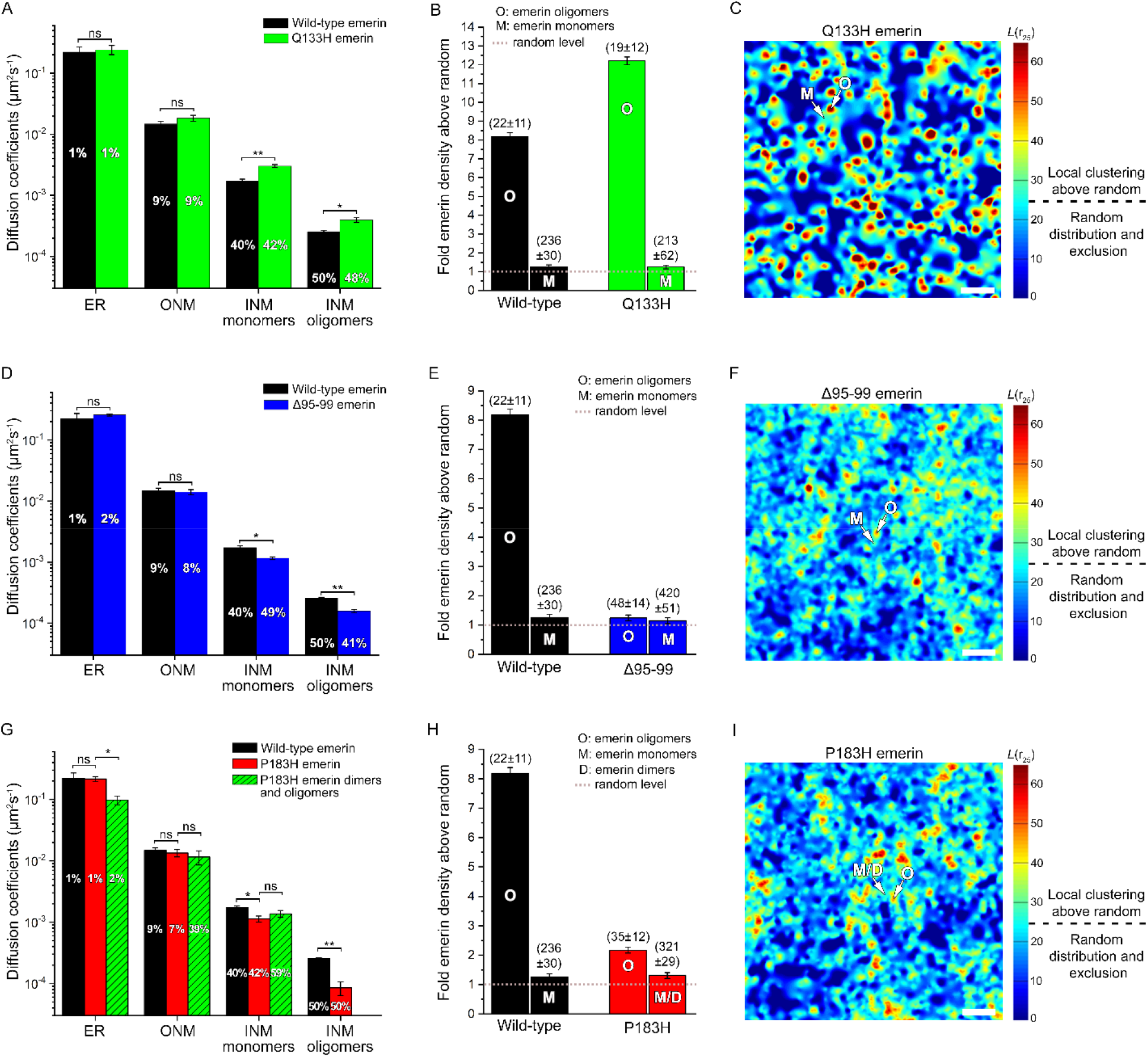
Emerin mutants exhibit modified lateral mobilities and defective oligomerization. (A) Diffusion coefficients (±s.e.m.) and percentages of wild-type and Q133H emerin. (B) Molecular densities above random (±s.e.m.) for wild-type and Q133H emerin oligomers (O) and monomers (M). (C) Local cluster map of Q133H emerin. M: monomer areas, O: oligomer nanodomains. Scale: 250 nm. (D) Diffusion coefficients (±s.e.m.) and percentages of wild-type and Δ95-99 emerin. (E) Molecular densities above random (±s.e.m.) for Δ95-99 (208092 localizations, 8 nuclei) and wild-type emerin oligomers (O) and monomers (M). (F) Local cluster map of Δ95-99 emerin. Scale: 250 nm. (G) Diffusion coefficients (±s.e.m.) and percentages of wild-type and P183H emerin compared to complemented P183H emerin-GFP-emerin species (H) Molecular densities above random (±s.e.m.) for P183H and wild-type emerin oligomers (O) and monomers/dimers (M/D). (I) Local cluster map of P183H emerin. M/D: monomer/dimer areas, O: oligomer nanodomains. Scale: 250 nm. For (A, D and G), t-test, ns: non-significant, *: *p*<0.05, **: *p*<0.01. For (B, E and H), values in parenthesis represent the size (±s.e.m.) of each domain in nanometers.

Although, it was reported that Q133H has a reduced capacity to self-assemble in vitro(Herrada et al., 2015), its spatial distribution and cluster maps show that it organizes into monomers and oligomers across the INM (Fig. 6B-C). Like wild-type emerin, Q133H monomers are distributed in 213±62 nm membrane domains (Fig. 6B), an indication that the inability to bind nuclear actin increases the mobility of Q133H monomers but does not affect their overall spatial distribution at the INM. Q133H also maintain the ability to self-assemble at the INM where it forms oligomeric clusters having sizes of 19±12 nm, comparable to the 22±11 nm size of wild-type emerin oligomers, but with molecular densities 12.2±0.2 fold above random (Fig. 6B-C; Table S2). This 50% increase in oligomerization compared to wild-type indicates that deficient binding of Q133H to nucleoskeletal actin leads to a disproportionate self-assembly of emerin into oligomeric nanodomains. It also implies that direct binding to nuclear actin normally reduces the oligomerization potential of wild-type emerin, consistent with our observation that excessive nuclear accumulation of actin impedes the formation of emerin oligomers at the INM (Fig. S7).

#### 95-99 mutation

When we performed similar studies with Δ95-99 emerin, we found that diffusions at the ER membrane and the ONM are unchanged compared to wild-type emerin (*p* not significant, Fig. 6D; Table S1), but that lateral mobilities at the INM are significantly reduced (*p*<0.05 and *p*<0.01, Fig. 6D), with the slowest Δ95-99 emerin population being essentially immobile. This suggests that Δ95-99 interacts more strongly or more frequently than wild-type emerin with some of its binding partners on the nucleoplasmic side of the NE. Previous biochemical studies have shown that the Δ95-99 deletion disrupts emerin interactions with most of its binding partners, including lamin A/C and actin, but not BAF(Berk et al., 2013b). As BAF binding strongly influences the mobility of wild-type emerin (Fig. 2A), the reduced diffusion of Δ95-99 at the INM could stem from repetitive interactions with BAF, as recently proposed(Samson et al., 2017). To destabilize these interactions, we attempted to track Δ95-99 in *EMD^−/y^* HDF knocked down for endogenous BAF and expressing BAF^L58R^. Co-expression of both Δ95-99 and BAF^L58R^, however, proved toxic to cells, indicating that interactions between Δ95-99 emerin and BAF are important to maintain cell viability.

In addition to its slow mobility, Δ95-99 displays defective oligomerization at the INM. Δ95-99 is distributed randomly over large, 420±51 nm NE domains and in smaller, 48±14 nm nanodomains where the molecular density of 1.3±0.1 fold above random is greatly reduced compared to wild-type emerin oligomers (Fig. 6E; Table S2). Effectively, Δ95-99 forms less dense and fewer oligomerization nanodomains in cluster maps (Fig. 6F). Thus, Δ95-99 does not efficiently oligomerize at the INM, consistent with its impaired self-assembly in vitro and reports that the deletion lowers emerin/emerin proximity(Herrada et al., 2015). Interestingly, despite an intact lamina and the preserved ability of Δ95-99 to bind SUN1(Haque et al., 2010), the molecular densities and cluster maps of Δ95-99 are similar to those of wild-type emerin when the expression of lamin A/C is reduced or when SUN1 LlNC complexes are disrupted (Fig. 3C-D). This reduced oligomerization of Δ95-99 to levels seen after lamin A/C depletion implies that direct binding to lamin A/C is required to stabilize emerin oligomers at SUN1 LINC complexes. Such a stabilizing role of lamin A/C is consistent with prior proximity ligation assays where Δ95-99 was found less close to lamin A/C than wild-type emerin(Herrada et al., 2015) and studies showing that the deletion abolishes direct lamin A/C binding to emerin(Lee et al., 2001). Together, these results show that Δ95-99 primarily distributes at random across the INM due to its reduced self-assembly, its inability to directly bind lamin A/C and its slow mobility.

#### P183H mutation

Like for the other emerin mutants, the diffusion of P183H emerin at the ER membrane and the ONM is unchanged compared to wild-type emerin (*p* not significant, Fig. 6G; Table S1). However, its INM mobility is reduced for the populations attributed to monomers and oligomers (*p*<0.05 and *p*<0.01, Fig. 6G), with the slowest P183H population being immobile. This slow diffusion resembles that observed for Δ95-99 and again implies that P183H interacts more frequently than wild-type emerin with some of its nucleoplasmic binding partners. In vitro, P183H binds SUN1(Haque et al., 2010), BAF and actin(Berk et al., 2013b) and displays enhanced binding to lamin A/C compared to wild-type emerin(Lee et al., 2001). Its reduced INM mobility might therefore be linked to an increased binding frequency to BAF, like for Δ95-99, or to its enhanced binding to lamin A/C. The slow diffusion of P183H could also be due to the formation of dimers as suggested based on its strong propensity to self-assemble in vitro(Herrada et al., 2015) and observations that residue P183 is positioned in the 168-186 emerin region required to limit emerin-emerin association(Berk et al., 2014). To determine if P183H forms dimers, we performed tracking by CALM after co-expression of P183H fused to split-GFP fragments. Three populations of P183H-GFP-P183H emerin species were detected: a 2% population with a mobility slightly slower than P183H at the ER membrane (*p*<0.05, Fig. 6G), a larger 39% population with a mobility comparable to the ONM behavior of P183H (*p* not significant, Fig. 6G), and a dominant 59% population with a lateral diffusion similar to the INM population initially attributed to P183H monomers by sptPALM (*p* not significant, Fig. 6G; Table S1). Surprisingly, no fluorescent species with diffusion coefficient matching that of the immobile P183H oligomers are detected (Fig. 6G). The high frequency detection of complemented P183H-GFP-P183H emerin species at both the ER membrane and the ONM indicates that P183H is indeed more prone to form dimers than wild-type emerin before reaching the INM. This apparent biased monomer:dimer equilibrium towards dimers is maintained at the INM where it precludes an efficient association of P183H into oligomer domains, in particular if dimers are stabilized by irreversible assembly of the split-GFP fragments. This indicates that P183H has a strong propensity to form dimers that could impact oligomerization at the INM. Consistent with these observations, the dimerization of P183H leads to a significantly reduced oligomerization at the INM. P183H monomers/dimers are distributed in domains with typical sizes of 321±29 nm and molecular densities of 1.3±0.1 fold above random (Fig. 6H; Table S2). Smaller, 35±12 nm nanodomains are also observed, but their molecular density is reduced to 2.2±0.1 compared to wild-type oligomers (Fig. 6H). Indeed, while P183H can still bind lamin A/C and SUN1, it forms oligomeric domains having lower molecular density than wild-type emerin in cluster maps (Fig. 6J), indicating that the dimerization of P183H hinders further self-association into dense oligomers at the INM.

### Abnormal reorganization of Δ95-99 emerin in response to mechanical stress

To characterize the importance of emerin oligomerization for nuclear deformation and responses to mechanical stress, we also studied the nanoscale reorganization of Δ95-99 in micropatterned cells. For deformed nuclei in 15 μm patterns, Δ95-99 monomers are dispersed over large, 810±215 nm INM domains, almost double the size of monomer domains in non-stressed nuclei (Fig. 7A). However, as stress becomes more pronounced in 10 μm patterns, the dispersion of Δ95-99 monomers recedes, with distributions in 499±250 nm domains (Fig. 7A). Thus, contrary to wild-type emerin, a progressive dispersion of Δ95-99 monomers over increasingly large INM areas is not observed with growing mechanical stress, consistent with the abnormal changes in NSI induced by the deletion. The formation of emerin oligomers remains very limited, with Δ95-99 molecular densities increasing slightly from 1.3±0.1 to 1.7±0.1 and 2.0±0.1 fold above random in 15 μm and 10 μm micropatterns (Fig. 7A; Table S2). With the concurrent enlargement of nanodomains from 48±14 nm to 81±16 nm and 75±20 nm, the relative oligomerization of Δ95-99 compared to non-mechanically stressed nuclei increases by 3.9 fold initially, but does not rise further as nuclear stress intensifies (Fig. 7B). Δ95-99 oligomeric nanodomains remain sparser and significantly less dense than for wild-type emerin in cluster maps of nuclei under stress (Fig. 7C). Thus, the failure of Δ95-99 to gradually self-associate at sufficiently high molecular densities leads to defective nuclear responses to force. This underlines the importance of modulating the oligomerization of emerin as a function of mechanical stress intensity for adaptive nuclear deformations.

**Fig. 7.**
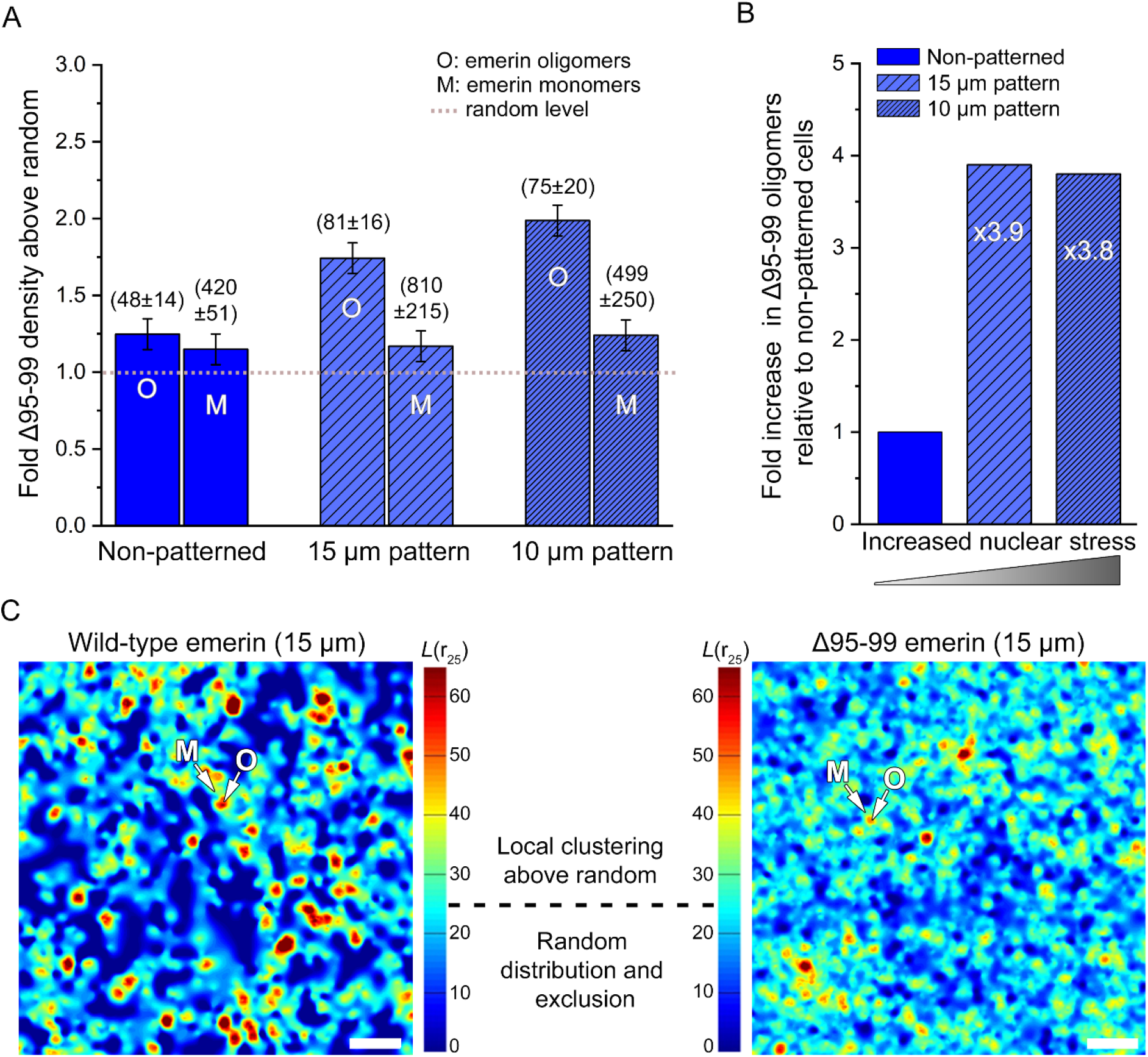
Insufficient oligomerization of Δ95-99 emerin mutant against mechanical stress. (A) Molecular densities above random (±s.e.m.) for Δ95-99 emerin oligomers (O) and monomers (M) in non-patterned cells or after nuclear deformation on 15 and 10 μm micropatterns. Values in parenthesis represent the size (±s.e.m.) of each domain in nanometers. (B) Changes in Δ95-99 emerin oligomers as a function of nuclear stress on 15 and 10 μm micropatterns. (C) Local cluster maps of wild-type and Δ95-99 emerin after nuclear deformation on 15 μm micropatterns M: monomer areas, O: oligomer nanodomains. Scale: 250 nm.

## Discussion

Combining single molecule imaging and quantitative measurements, we showed that emerin distributes as monomers and oligomers at the INM and that clinically relevant EDMD mutations induce modified emerin mobility and nanoscale organizations. The oligomerization of emerin is modulated by its ability to engage or disengage interactions with various structural elements juxtaposed to the INM, including lamin A/C, SUN1, nuclear actin and BAF. Regulated expressions of lamin A/C and SUN1, together with functional LINC complexes, are essential for the formation and the stabilization of emerin oligomers, while balanced interactions of emerin with BAF and nuclear actin further modulate its diffusion and its oligomerization potential. We also showed that the mechanotransducing functions of emerin are coupled to its oligomeric state, with the formation and maintenance of emerin oligomers being central to nuclear shape adaptation against forces. Indeed, EDMD-inducing mutations that affect the self-assembly of emerin and its binding to structural elements at the INM lead to abnormal nuclear deformations and defective nucleus positioning in response to mechanical stress.

While the altered in vitro binding properties of Q133H, Δ95-99 and P183H emerin mutants result in expected differences in organization in cells, they do not necessarily induce fully predicable changes in diffusion and distribution at the NE. In effect, via its flexible IDR, emerin appears to mediate complex interactions with itself, lamin A/C, SUN1, nuclear actin and BAF, where binding to one partner impacts oligomerization and, interdependently, affects interactions with other partners. As shown from the organization of wild-type emerin after nuclear actin depletion or accumulation and our study of Q133H emerin, binding to nuclear actin significantly influences the mobility and the oligomerization potential of emerin. Actin binding to the IDR of emerin could mask interaction domains that normally serve as LEM binding and self-association sites between emerin monomers(Berk et al., 2014). IDR masking, combined with the reduced lateral mobility of actin-bound emerin, might therefore modulate the formation of oligomers by limiting molecular collisions between emerin monomers. Consistent with the need to precisely adjust the oligomerization potential of emerin for adaptive nuclear shape deformations, increased emerin diffusion and controlled formation of emerin oligomers are coupled molecular events during nuclear adaptation against stress, as shown by our measurements in micropatterns. We thus speculate that the deficient binding of Q133H emerin to nuclear actin and its over-oligomerization at SUN1 LINC complexes are linked to the abnormal nuclear deformations observed in cells.

The differential interactions of emerin with its nucleoplasmic partners and their influence on its mobility and its oligomerization are also underlined by the surprisingly slow mobility of Δ95-99 emerin, which does not bind lamin A/C nor actin in vitro, but binds BAF. The 48-118 region of emerin, where Δ95-99 is located, was suggested to act as a binding site for the LEM domain(Berk et al., 2014) and through altered region flexibility, the deletion might reduce the efficacy of such interactions, potentially causing the LEM domain to bind BAF more frequently(Samson et al., 2017). Repeated interactions of Δ95-99 with BAF and, indirectly, with the lamina or chromatin via ternary BAF complexes, could slow its diffusion, limit molecular collisions, and impede its oligomerization by sequestering the LEM domain and reducing bridging interactions with LEM binding sites on other emerin. The Δ95-99 deletion, within the 55-132 lamin tail-binding region of emerin(Berk et al., 2014), could also prevent a stabilization of already sparse Δ95-99 oligomers by direct lamin A/C binding. We thus postulate that the aberrant nuclear deformations against stress of cells expressing Δ95-99 emerin stem from out of balance interactions of this mutant with BAF and its inability to form lamin A/C-stabilized oligomers at SUN1 LINC complexes.

A similar impediment of *in trans* interactions between the LEM domain and self-association sites along the joined IDR of P183H dimers likely result in their reduced ability to oligomerize, despite retained binding to lamin A/C and SUN1. The reported enhanced binding of lamin A/C to P183H emerin(Lee et al., 2001), in close proximity to these same self-association sites could also interfere with inter-emerin bridging interactions. In both cases, the inability of the LEM domain to access binding sites along the IDR would elicit its repeated interactions with BAF, leading to the observed slow diffusion of P183H at the INM. The defective nuclear shape adaptation to mechanical challenges of cells expressing P183H might thus arise from emerin dimerization and excessive interactions with BAF or lamin A/C that prevent an efficient formation of emerin oligomers at LINC complexes. Our observation that lamin A/C overexpression, which was previously shown to induce the nuclear accumulation of BAF(Loi et al., 2016), leads to the decreased oligomerization of wild-type emerin is indeed consistent with the need to modulate emerin/BAF/lamin A/C tripartite interactions to promote the self-assembly of emerin.

As we have shown, because Δ95-99 emerin does not self-assemble into sufficiently dense oligomers at the INM, it induces aberrant nuclear shape remodeling against mechanical stress. The phosphorylation of emerin residues Tyr 74 and Tyr 95 was previously shown to mediate the recruitment of lamin A/C to the LINC complex during nuclear stiffening in response to force(Guilluy et al., 2014). Our observation that Δ95-99 fails at promoting adaptive nuclear deformation in response to mechanical cues by its reduced ability to form lamin A/C- and SUN1-stabilized oligomers is consistent with the major role played by Tyr 95 phosphorylation for emerin-mediated mechanotransduction at the NE. The abnormal organization of cytoskeletal actin in mechanically challenged cells expressing Δ95-99, akin to disorganizations detected with a non-phosphorylable tyrosine 74-95FF emerin(Guilluy et al., 2014), also suggests that emerin oligomers strengthen the connection between lamin A/C, the LINC complex and cytoskeletal filaments to promote correct nucleus positioning and deformation. These data point towards a link between Tyr 95 phosphorylation, emerin oligomerization and recruitment of lamin A/C for stiffening the NE at LINC complexes, likely driven by phosphorylation-induced changes in emerin conformation(Berk et al., 2013a; Tifft et al., 2009). It is thus possible that emerin oligomeric nanodomains are enriched in Tyr 95 phosphorylated emerin, lamin A/C and LINC complex components, while emerin monomers populate the rest of the INM, consistent with emerin enrichments in distinct nucleoskeletal “niches” at the NE(Berk et al., 2013a).

Taken together, these lines of evidence support a model in which, during stress responses, transient unbinding of emerin monomers from nuclear actin and BAF favors oligomerization by increasing the lateral diffusion of emerin at the INM and exposing both its LEM domain and self-association sites along its IDR for intermolecular binding between emerin at LINC complexes (Fig. 8). Within emerin oligomers, stabilization by direct interactions with lamin A/C and SUN1 and additional modulation of emerin self-association by nuclear actin and BAF likely allow for a precise regulation of the oligomerization potential and the size of emerin oligomeric nanodomains at LINC complexes, as required for adaptive nuclear deformation in response to forces transmitted by cytoskeletal filaments. The localized INM distribution of emerin oligomers, the involvement of lamin A/C and SUN1 in their stabilization, their specific disruption following LINC complex destabilization, and their reliance on a part of the emerin IDR that requires phosphorylation for the recruitment of lamin A/C to LINC complexes(Guilluy et al., 2014), indeed support the idea that they are sites where interactions between emerin, lamin A/C and LINC components are strengthened. It was recently suggested that the clustering of SUN proteins could enable them to sustain high mechanical loads at the NE during force-dependent processes(Jahed et al., 2018). Emerin oligomerization at SUN1 LINC complexes, might therefore contribute to the increased connectivity between the nucleoskeleton, the NE and the cytoskeleton for anchoring the nucleus and providing force absorption contact points during nuclear deformation.

**Fig. 8.**
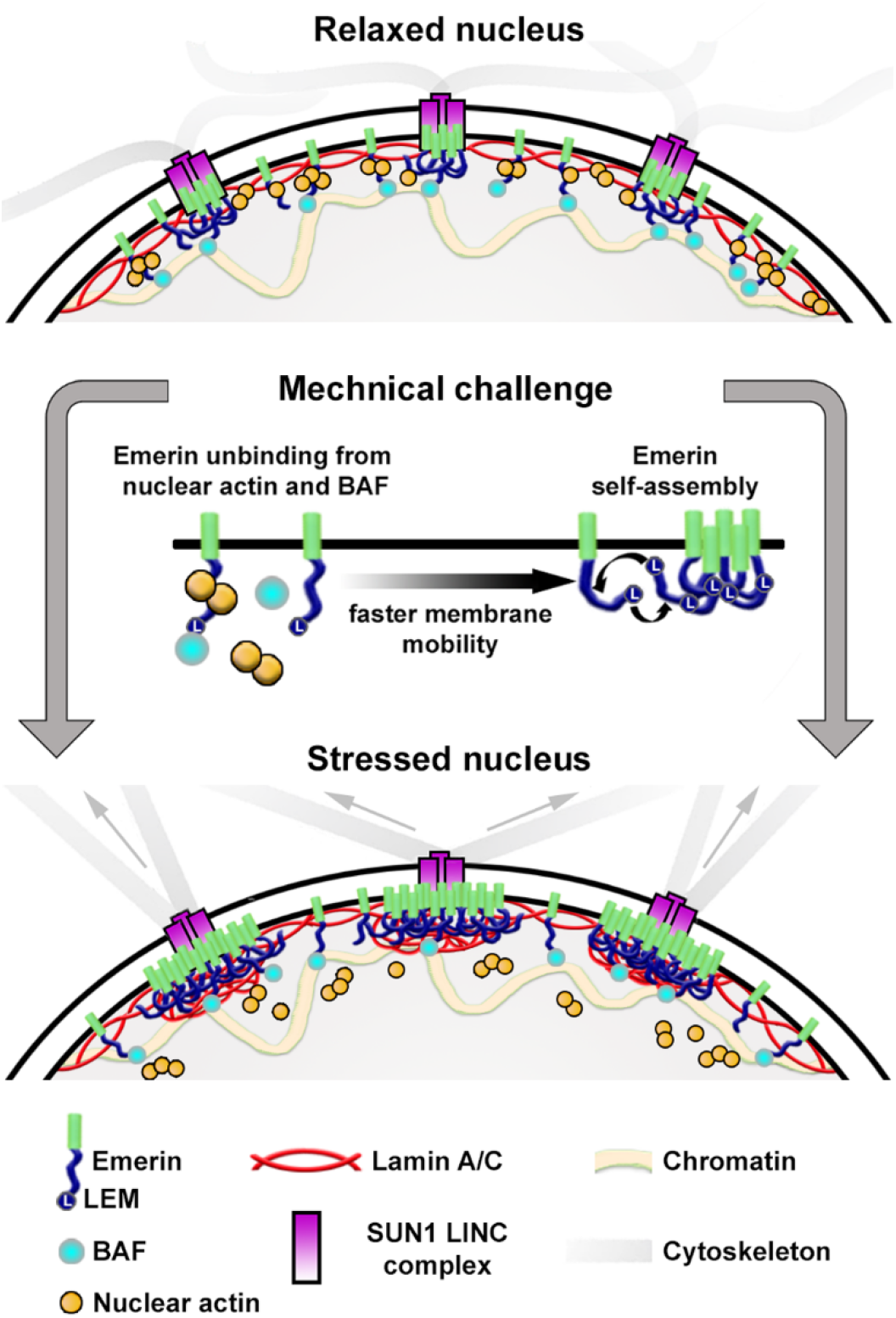
Model of emerin re-organization at the nuclear envelope in response to mechanical challenges. Emerin monomer unbinding from nuclear actin and BAF induces increased lateral mobility at the inner nuclear membrane and favors LEM domain interactions with binding sites along the intrinsically disordered region of other emerin, for the controlled formation of emerin oligomers at SUN1 LINC complexes and their stabilization by lamin A/C.

In response to surging mechanical stress, the incremental oligomerization of emerin in enlarged nanodomains we observed could be part of a mechanism that redistributes increasing forces over wider areas at LINC complexes to maintain a basal membrane pressure at these anchoring points between the nucleoskeleton, the NE and the cytoskeleton. Beyond this central role played by emerin self-assemblies, emerin might have additional functions outside these anchoring oligomeric nanodomains. For instance, the disengagement of emerin monomers from nuclear actin likely reduces NE contacts with the nucleoskeleton, while some membrane connectivity with nuclear chromatin is maintained, via BAF. Indeed, both the nucleoskeleton and chromatin tethering to the NE participate in nuclear mechanics(Stephens et al., 2017). Together, strengthening the connections between the NE, the lamina and the cytoskeleton at LINC complexes, but partially relaxing them in the rest of the membrane could provide means to couple controlled nuclear deformation with nucleus positioning in cells. As we showed, such coupling is defective with emerin mutants.

Additional cross-correlative studies will be needed to further define how the structural interdependencies between emerin monomers, emerin oligomers, SUN1, BAF, and key nucleoskeletal elements spatially regulate INM/lamina and INM/chromatin contacts and reinforce the connections between the NE and the cytoskeleton at LINC complexes during nuclear responses to force. At the center of these processes, emerin diffusion and monomer/oligomer exchanges provide a means to transduced mechanical cues throughout the entire NE, for coordinated changes in local nuclear stiffness and remodeling of the nuclear shape.

## Materials and Methods

### Cell culture, emerin expression and cell staining

Emerin-null human dermal fibroblast (*EMD*^−/y^ HDF) and normal dermal fibroblasts (*EMD*^+/y^ HDF) were kindly provided by Dr. Howard Worman, Columbia University, USA, and regularly tested for contaminations. *EMD*^−/y^ HDF are derived from a male EDMD patient (G-9054) and carry a 59 nucleotide deletion within the *EMD* gene (*EMD* g.329del59) that ablates emerin expression(Talkop et al., 2002). HDF were grown in DMEM (Lonza) with 10% fetal bovine serum (Gibco-Life Technologies), 50 units ml^−1^ penicillin and 50 μg ml^−1^ streptomycin and maintained at 37°C in a humidified atmosphere with 5% CO_2_.

Human wild-type emerin cDNA was kindly provided by Dr. Juliet Ellis, University College London, UK. For the expression of PA-TagRFP-emerin, a pEGFP-N1 plasmid backbone encoding emerin fused to the C-terminus of PA-TagRFP was produced by XbaI and KpnI insertion and PCR fusion of the human emerin cDNA. Cells plated on fibronectin-coated glass coverslips were transfected with PA-TagRFP-emerin using X-tremeGENE HP (Roche). 48-72 hours post-transfection, live cells were imaged by sptPALM in HBSS buffer at 37°C. For micropatterning experiments, cells grown on 6-well plates were trypsinized after 48-72 hours of transfection and plated on fibronectin-micropatterned coverslips.

To express SNAP-emerin, human emerin was first fused to the C-terminus of a SNAP tag by AscI and XhoI insertion in a pSNAP-tag(m) plasmid (NEB). SNAP-emerin was then subcloned into a modified pFUW lentiviral vector by NheI and AgeI insertion. Lentiviral particles for the expression SNAP-emerin were produced by the UCLA Vector Core. Transduction of HDF grown at 70% confluence on 6-well plates was done for 48 hours, using 25 ng ml^−1^ of lentiviral particles in complete growth medium containing 8 μg ml^−1^ of polybrene, after which the medium was replaced. Following another 24 hours incubation, cells were trypsinized, and plated on fibronectin-coated or fibronectin-micropatterned coverslips. For imaging, cells were fixed with 4% paraformaldehyde in PBS for 15 min, permeabilized with 0.1% Triton X-100 (Sigma-Aldrich) for 15 min and blocked with 4% bovine serum albumin (BSA, Sigma-Aldrich) + 0.1% Tween-20 (Sigma-Aldrich) for 30 min, at room temperature. Cells were then stained with 1 µM of SNAP-Surface-AlexaFluor 647 (BG-A647, NEB) in 4% BSA + 0.1% Tween-20 for 1 hour at 37°C, then thoroughly washed before super-resolution imaging.

For the expression of sGFP-emerin, humanized cDNA for split-GFP 1-10(Pinaud and Dahan, 2011) was inserted by NheI and XbaI digestion in the pEGFP-N1 plasmid backbone encoding emerin, and expressed as an N-terminal fusion to emerin. The shorter 11^th^ β-sheet M3 fragment(Pinaud and Dahan, 2011) was also fused to the N-terminus of emerin by PCR cloning using primers encoding the M3 fragment sequence and subcloning into the pEGFP-N1 backbone via NheI and XhoI digestions. Both plasmids were co-transfected in HDF using X-tremeGENE HP (Roche) as described for PA-TagRFP-emerin, and cells were imaged by CALM 48-72 hours post-transfection. All constructs were verified by sequencing.

For immunostaining of emerin, cells were grown on coverslips, fixed and permeabilized as described for SNAP-emerin staining. Cells were labeled with a rabbit anti-emerin antibody (1:500, Santa Cruz Biotechnology, sc-15378) for 45 min, washed in PBS and further labeled with a goat anti-rabbit Alexa fluor 488 antibody (1:400, Invitrogen) for 45 min. After washing, cells were mounted in a DAPI-Fluoromount G (Electron Microscopy Sciences) and imaged by confocal microscopy.

To label cytoplasmic actin and measure nuclear shape indices, cells were fixed with 4% paraformaldehyde in PBS for 15 min, permeabilized with 0.1% Triton X-100 for 10 min, and blocked with 4% bovine serum albumin + 0.1% Tween-20 for 1 hour. Cells were stained with phalloidin-iFluor 488 (1:1000, Abcam) for 1 hour, washed with PBS, mounted in DAPI-Fluoromount G and imaged by confocal or wide field microscopy.

### Mutations, siRNA and exogenous protein expressions

Emerin mutations Q133H and P183H were introduced in PA-TagRFP-emerin, SNAP-emerin, and emerin fused to both split-GFP fragments by site-directed mutagenesis using QuickChange Lightning Site Directed Mutagenesis (Agilent) and mutagenic primer pairs for:

Q133H: 5’-CGCTTTCCATCACCATGTGCATGATGA-3’ and 5’-GATCGTCATCATGCACATGGTGATGGA-3’

P183H: 5’-CCTGTCCTATTATCATACTTCCTCCTC-3’ and 5’-GTGGAGGAGGAAGTATGATAATAGGA-3’.

The Δ95-99 emerin deletion was produced using partially phosphorothioated PCR primers and T7 exonuclease digestion, as previously described(Stoynova et al., 2004). Primers pairs for the Δ95-99 deletion were:

5’-GACTACTTCACCA*C*C*A*GGACTTAT-3’ and
5’-GGTGAAGTAGTCG*T*C*A*TTGTAGCC-3’,

where * denotes phosphorothioate modifications. All primers were obtained from Integrated DNA Technologies (IDT) and all mutations were verified by sequencing.

siRNA duplex for BAF, SUN1 and Dicer siRNA for lamin A/C, SUN1 and controls were obtained from IDT. The sequences of sense nucleotides were as follows:

- BAF siRNA: 5’-AGAUUGCUAUUGUCGUACUUU-3’
- Lamin A/C DsiRNA #1: 5’-AGCUGAAAGCGCGCAAUACCAAGaa-3’,
- Lamin A/C DsiRNA #2: 5’-GGAACUGGACUUCCAGAAGAACAtc-3’,
- SUN1 DsiRNA #1 5’-GCUUUUAGUAUCAACCACGUGUCaa-3’.
- SUN1 siRNA #2: 5’-CCAUCCUGAGUAUACCUGUCUGUAU-3’.
- Control DsiRNA: 5’-CGUUAAUCGCGUAUAAUACGCGUat-3’ (IDT#51-01-14-04).

siRNA duplex for IPO9 was obtained from Ambion (id# S31299) and siRNA for the nuclear actin exporter XPO6 was obtained from Qiagen (id# SI00764099). All siRNA were transfected or co-transfected with emerin plasmids at 25 nM using X-tremeGENE HP (Roche). When associated with lentiviral expression of emerin, siRNA transfection was done 2 hours before viral titer application.

Exogenous BAF^L58R^ was expressed from an EGFP-BAF^L58R^ lentiviral plasmid(Samwer et al., 2017) and lentiviral particles were produced by the UCLA Vector Core. HDF cells were transduced with 25 ng ml^−1^ of lentiviral particles as described for SNAP-emerin. For rescue of lamin A/C expression and overexpression experiments, cDNA encoding for an exogenous siRNA-resistant human lamin A/C (Frock et al., 2006) was kindly provided by Dr. Richard Frock, Stanford University, USA and used together with lamin A/C DsiRNA #2, designed to knockdown endogenous lamin A/C. For rescue of SUN1 expression and overexpression experiments, cDNA encoding for an exogenous siRNA-resistant human SUN1 fusion to EGFP (EGFP-SUN1) was kindly provided by Dr. Christine Doucet, Centre de Biochimie Structurale, Montpellier, France and used together SUN1 siRNA #2, designed to knockdown endogenous SUN1, as previously described(Talamas and Hetzer, 2011). Exogenous expression of mcherry-DN-KASH was achieved by cell transfection of the mammalian expression plasmid #125553, obtained from Addgene.

### Cell micropatterning and nuclear shape index measurements

HDF were micropatterned as described previously(Bautista et al., 2019; Fernandez et al., 2017). Briefly, hexamethyldisilazane-activated glass coverslips (Marienfeld, #1.5, Ø25 mm) were stamped with rectangular and fibronectin-coated polydimethylsiloxane stamps having lengths of 210 µm and widths of 15 µm, 10 µm, or 5 µm respectively. Cell attachment outside the fibronectin strips was blocked with a 1% solution of Pluronic F-127. After attachment for 1 hour and removal of unattached cells, HDF were allowed to spread out on the micropatterns for 6 hours at 37°C before being prepared for microscopy imaging.

Using ImageJ(Schneider et al., 2012), the nuclear shape index (NSI) was determined by measuring the nuclear cross-sectional area and the nuclear perimeter of DAPI-stained nuclei imaged by wide-field microscopy, and by calculating the ratio:

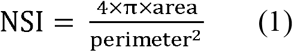

The NSI measures the roundness of the nucleus, with an NSI of 1 corresponding to a circular nuclear shape. Mean NSI values ± standard deviation of the mean were reported for multiple cell nuclei per condition. In graphs, the box length indicates the NSI interquartile range, the central square indicates the NSI mean, the central bar indicates the NSI median and the error bars indicate the standard deviation. The number of measured nuclei was n=54, 70, 61 and 62 nuclei for *EMD^+/y^* cells in non-patterned, 15, 10 and 5 µm wide patterns, respectively; n=75, 60, 70 and 66 nuclei for lamin A/C depleted cells; and n=62, 71, 66 and 46 nuclei for nuclear actin depleted cells. For *EMD^−/y^* cell expressing wild-type emerin, the number of measured nuclei was n=38, 33, 26 and 57 nuclei in non-patterned, 15, 10 and 5 µm wide patterns, respectively; n= 74, 58 and 37 nuclei for Q133H emerin; n= 64, 89, 45 and 46 nuclei for Δ95-99 emerin and n=82, 78 and 28 nuclei for P183H emerin. Significant differences between NSI values were evaluated using a Wilcoxon rank-sum test.

### Microscopy imaging

Confocal imaging of immunostained emerin in normal HDF, emerin-null HDF and emerin null HDF after expression of wild-type PA-TagRFP-emerin were performed on an Olympus Fluoview FV1000 confocal microscope equipped with a 60x/1.40 NA objective, 405 nm and 488 nm lasers and appropriate emission filters for imaging DAPI (450/20 nm) and Alexa-488 labeled antibodies against emerin (510/10 nm).

Confocal imaging of nuclear deformation and actin organization in micropatterned *EMD^+/y^* HDF was performed on a Zeiss LSM 700 microscope equipped with a C-Apochromat 63×/1.15 W Korr objective, excitation lasers at 405 nm and 488 nm and appropriate beamsplitter and emission channel settings for dual detection of DAPI and phalloidin-iFluor 488.

Wide-field imaging of labeled actin and labeled nuclei for NSI measurements were performed on an inverted Nikon Eclipse Ti-E microscope equipped with a 40x objective (Nikon), an iXon Ultra EMCCD camera (Andor), 405 nm and 488 nm lasers, a T495lpxr dichroic mirror and a 525/50 emission filter (Chroma) for phalloidin-iFluor 488 or a 458Di02 dichroic mirror and a 483/32 emission filter (Semrock) for DAPI.

SptPALM, dSTORM and CALM imaging were performed on an inverted Nikon Eclipse Ti-E microscope equipped with a 100x 1.49 NA objective (Nikon), an iXon EMCCD camera (Andor), perfect focus drift compensation optics, an astigmatic lens for 3D super-resolution imaging, a piezo z-scanner for calibration of 3D super-resolution images (Mad City Labs), laser lines at 405, 488, 561 and 647 nm (Agilent), a multiband pass ZET405/488/561/647x excitation filter (Chroma), a quad-band ZT405/488/561/647 dichroic mirror (Chroma) and appropriate emission filters for sptPALM imaging of PA-tagRFP (600/50 nm, Chroma), 3D-dSTORM imaging of Alexa-647 (700/75 nm, Chroma) and CALM imaging of complemented split-GFP (525/50 nm, Semrock).

sptPALM of PA-TagRFP-emerin was performed in 37°C HBSS buffer (Corning) by highly inclined and laminated optical sheet (HILO) excitation of the bottom nuclear membrane of cells with a continuous and low power photoactivation at 405 nm and an excitation at 561 nm. The HILO illumination angle was θ= 51.6°. Images were acquired continuously at a frame rate of 40 ms per frame for no longer than 3 minutes per cell to limit UV damage. CALM imaging of complemented emerin-GFP-emerin species was done as described for sptPALM but with a single HILO excitation at 488 nm.

dSTORM of SNAP-emerin labeled with BG-A647 was performed at room temperature in a photoswitching buffer composed of 10% glucose, 0.5 mg ml^−1^ glucose oxidase (Sigma), 40 μg ml^−1^ catalase (Sigma), and 1% β-mercaptoethanol (Sigma). Continuous photoswitching was achieved with a low power 488 nm laser and imaging was done with a 647 nm laser excitation at a frame rate of 80 ms per frame. Z-calibration and sample drift corrections were done using a few 40 nm TransFluoSphere beads (488/685 nm, Life Technologies) as fiducial markers spread on the cell samples.

For rescue and overexpression experiments, dSTORM imaging of SNAP-emerin was performed only in cells expressing exogenous human lamin A/C, human EGFP-SUN1 or mcherry-DN-KASH, as identified by lamin A/C immunostaining, EGFP or mCherry fluorescence. In the case of lamin A/C and EGFP-SUN1 rescues, imaging was done in cells having relatively low exogenous expressions.

### Fluorescence recovery after photobleaching (FRAP) of emerin

FRAP of wild-type emerin and emerin mutants was done in U2OS cells transiently transfected with PA-TagRFP-emerin. Cells were imaged at 37°C in HBSS buffer (Corning) on an Olympus Fluoview FV1000 confocal microscope equipped with a 60x/1.40 NA oil immersion objective. PA-TagRFP-emerin at the bottom NE was briefly photoactivated by laser scanning at 405 nm and PA-TagRFP fluorescence was monitored every 2.8 seconds with a 543 nm laser. Circular regions of interest (7 µm in diameter) were photobleached and recorded. After background subtraction, FRAP curves were doubly normalized as previously described(Phair et al., 2004) to correct for loss of fluorescence due to bleaching during acquisition. FRAP curves from measurements on 9-12 cells per condition were averaged after normalization to full scale. After comparison of one- or two-component lateral diffusion models(Soumpasis, 1983), the apparent diffusion coefficients of emerin were determined by fitting FRAP curves with a two-component model using equations for a uniform circular bleach region(Soumpasis, 1983). A fast (D_fast_ = 4.48×10^−2^ ± 1.0×10^−2^ μm^2^/s, 17%) and slow (D_slow_ = 4.0×10^−3^ ± 4×10^−4^ μm^2^/s, 83%) diffusive behaviors of wild-type emerin is observed by FRAP (Fig. S1), consistent with previously diffusion values (1.0×10^−1^ ± 1×10^−2^ m^2^/s and 7.50×10^−2^ ± 2.9×10^−3^ m^2^/s)(Haraguchi et al., 2008; Ostlund et al., 1999; Ostlund et al., 2006) and the reported presence of a nearly immobile emerin fraction at the NE(Haraguchi et al., 2008; Shimi et al., 2004). Emerin mutations P183H, Q133H, and Δ95-99 do not display statistically significant differences in ensemble diffusion dynamics compared to wild-type emerin (Fig. S1).

#### Cell extracts and immunoblotting

After siRNA treatment, cells were harvested and fractionated as described previously(Berk et al., 2013a). Cells were scraped, centrifuged for 5 min at 4000 g and washed three times with PBS. Cell pellets were then flash frozen and stored at −80°C. Cells were thawed for 10 minutes on ice, and then for 10 min in 300 µl ice cold hypotonic lysis buffer (20 mM HEPES, pH 7.4, 50 mM N-acetylglucosamine, 1 mM DTT, 100 µM PMSF, 1 µg/ml pepstatin A, 1x protease inhibitor (Roche) and 1x PhosSTOP phosphatase inhibitor (Roche) before being resuspended and set on ice for another 10 min. Cells were then spun down at 17,000 g for 1 min to collect the supernatant cytoplasmic fraction. The pellet was then washed three times in PBS with spinning at 17,000 g for 1 min in between washes to remove any residual cytoplasmic components. The pellet was then resuspended in nuclear lysis buffer (50 mM Tris-HCL, pH 7.4, 300 mM NaCl, 0.3% Triton X-100, 50 mM N-acetylglucosamine, 1 mM DTT, 100 µM PMSF, 1 µg/ml pepstain A, 1x protease inhibitor (Roche) and 1x PhosSTOP phosphatase inhibitor (Roche) before vortexing to release nuclear contents. Nuclear fractions were additionally sonicated 20 times in 0.5 s bursts to liberate dense nuclear aggregates. Cytoplasmic and nuclear fractions were analyzed by SDS-PAGE. Protein concentrations were determined using a Bradord assay, and equal amounts of protein were loaded on gels before running in a Laemmli Buffer. Proteins were transferred to a nitrocellulose membrane (Bio-Rad) by wet transfer at 4°C. The membrane was then rinsed with Tris-buffered saline (TBS, pH 7.5) and blocked with 5% milk in TBS for 1 hour. Membranes were probed with the following primary antibodies: mouse anti-lamin A/C (1:1000, Santa Cruz Biotechnology, sc-7292), mouse anti-actin (1:1000, Santa Cruz Biotechnology, sc-8432), mouse anti-BAF (1:1000, Santa Cruz Biotechnology, sc-166324), rabbit anti-SUN1 (1:2000; HPA008346, Sigma-Aldrich), mouse anti-histone H2A (1:1000, Santa Cruz Biotechnology, sc-515808) and mouse anti-GAPDH (1:1000, GeneTex, GTX627408). A goat anti-mouse IgG (H+L) HRP conjugate (1:3000, Invitrogen, 62-6520) and a goat anti-rabbit IgG (H+L) HRP conjugate (1:5000, Invitrogen, 65-6120) were used as secondary antibodies. Blots were developed with SuperSignal West Femto Maximum Sensitivity Substrate (Thermo) and imaged on a Chemidoc XRS+ (BioRad). All assays were performed in triplicate and immunoblot quantification was done with ImageJ, using t-tests for statical comparisons with wild-type (Fig. S2).

#### Immunostaining and fluorescence imaging of RNAi effects and exogenous protein expressions

*EMD^+/y^* HDF were transfected with control siRNAs, or siRNAs against lamin A/C, IPO9, BAF, and SUN1, respectively as described in Materials and Methods. Cell were fixed with 4% PFA in 1x PBS for 15 min, permeabilized with 0.1% Triton X-100 for 10 min, and blocked with 4% BSA + 0.1% Tween-20 for 1h, all at RT. For lamin A/C and SUN1 immuno-staining, cells were incubated with mouse anti-lamin A/C (1:400; sc-7292, Santa Cruz Biotechnology) and rabbit anti-SUN1 (1:500; HPA008346, Sigma-Aldrich) for 1hr at RT, rinsed 3x with 1x PBS for 5 minutes each, then stained with goat anti-mouse-Alexa Fluor 647 (1:500, Life Technologies) and goat anti-rabbit-Alexa Fluor 488 (1:500, Life Technologies) for 1hr at RT. Following 3x PBS rinse, the coverslips were mounted in DAPI-Fluoromount G (Electron Microscopy Sciences) and imaged. For BAF staining, cells were incubated with mouse anti-BAF (1:100; sc-166324, Santa Cruz Biotechnology) for 2hr at RT, then with goat anti-mouse-Alexa Fluor 647 (1:500) for further 2hr. To assess the effect of BAF L58R re-expression on lamin A/C organization, 24 hours after siRNA treatment against BAF, cells were transduced with the lentivirus for EGFP-BAF L58R expression and fixed two days later, before staining for lamin A/C as described above. To assess the influence of lamin A/C or EGFP-SUN1 expression on the localization of endogenous emerin, *EMD^+/y^* HDF were fixed and immunostained with rabbit anti-emerin (Abcam) and mouse anti-lamin A/C (Santa Cruz Biotechnology) antibodies as described above. Confocal fluorescence imaging (Fig. S2 and S6) was performed on a Zeiss LSM 780 microscope equipped with a C-Apochromat 63×/1.15 W Korr objective, excitation lasers at 405 nm, 488 nm and 633 nm, and appropriate beam splitter and emission channel settings for detection of DAPI, GFP, Alexa Fluor-488 and Alexa Fluor-647 labeled secondary antibodies. Other than effective and specific knockdown of targeted protein expressions, there was no obvious indirect or off-target effects of the different siRNA treatments on the organization other studied proteins (Fig. S2).

To assess the influence of IPO9 and XPO6 siRNA on nuclear actin organization and content, Utr230-EGFP-3xNLS (Utr230-EN, Addgene #58466) was subcloned into a pHR-SFFV lentiviral plasmid backbone, viral particles were generated, and *EMD^+/y^* HDF cells were transduced with 25 ng ml-1 of lentiviral particles as described for SNAP-emerin. 10 µg/mL blasticidin S (Invivogen) was used over 12 days to establish a stable cell line expressing Utr230-EN. Cells were then transfected with 25 nM of control, IPO9 or XPO6 siRNAs, as described above, before being fixed with 4% PFA for 15 minutes, rinsed 3x with 1xPBS, and mounted using Fluoromount-G (Electron Microscopy Sciences). Confocal fluorescence imaging was performed on a Zeiss LSM 780 upright microscope equipped with a Plan-Apochromat 40×/1.4 Oil objective, excitation laser at 488 nm, and appropriate beam splitter and emission channel settings for the detection of EGFP. From confocal images, cells were classified into three groups depending on the nuclear localization pattern of Utr230-EN: (i) small puncta and diffuse, (ii) diffuse or (iii) large foci, based on a similar previous classification(Belin et al., 2013). Data were pooled from three independent assays for each condition.

#### Quantification of ONM emerin fraction

The fraction of ONM emerin in *EMD^+/y^* HDF was quantified after saponin (ONM+ER emerin) or Triton X-100 (INM+ONM+ER emerin) permeabilization and immunostaining with rabbit anti-emerin (1 µg/mL, ab40688, Abcam) and mouse anti-lamin A/C (1:1000, sc-7292, Santa Cruz Biotechnology) antibodies, as previously described(Bautista et al., 2019). Whole cell volumes were imaged by 3D confocal microscopy and all the z-slices were combined into a single image by sum intensity z-slices projection. Regions of interest were delineated to quantify emerin fluorescence intensities over the entire cells (ROI1) or over the nucleus (ROI2) after background corrections. Nuclear ROI2 were further corrected for ER emerin contributions by evaluating the mean ER-only emerin intensity per area from ROI1-ROI2 intensities. Using a representative set of 20 cells for both saponin and Triton X-100 treatments, the fraction of ONM emerin per cell was then calculated using:

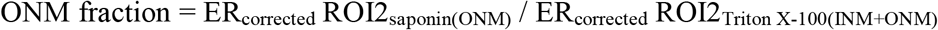

The mean ONM fraction of emerin at the NE was 13±6 %, in good agreement with the levels of ONM emerin detected in sptPALM experiments (Fig. S4).

### Analyses of diffusion coefficients

Localization and tracking analyses were performed using the software package SLIMfast, which uses multiple-target tracing algorithms(Serge et al., 2008) and was kindly provided by Christian Ritcher and Jacob Piehler. Localizations were performed by 2D-gaussian fitting of the point-spread-function of each activated PA-TagRFP-emerin or activated emerin-GFP-emerin species in each frame. Localization precision was determined as previously described(Mortensen et al., 2010; Thompson et al., 2002), and individual PA-TagRFP-emerin were localized with a precision of 13±5 nm. Diffusion trajectories were built by linking localizations frame to frame and accounting for blinking statistics and local particle densities. Trajectories with fewer than three steps were discarded. Diffusion coefficients were estimated using a probability density of square displacement (PDSD) analysis(Schutz et al., 1997). For each time lag *t*, the PDSD curve was fitted with the following model:

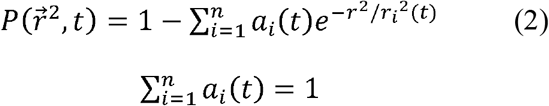

where *r_i_ (t)* is the square displacement and *a_i_(t)* is the population density of *i* numbers of diffusive behaviors at each time lag *t*. To limit the risks of overfitting or underfitting the PDSD curves and select an appropriate model for *i* numbers of diffusive behaviors in each data set, we used both an Akaike information criterion (AIC) and a Bayesian information criterion (BIC) after fitting PDSD with models where 1≤ *i* ≥5. Square displacement curves (*ri^2^(t)*) were extracted from PDSD analyses and reported with error bars determined using 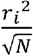 where N is the number of analyzed trajectories per time lag, as previously described(Schutz et al., 1997). Diffusion coefficients (*D*) representative of each behavior were determined by fitting each *ri^2^(t)* curves over the first four time lags (*t*_1_-*t*_4_) using OriginPro 2020 software (OriginLab) and a 2D Brownian diffusion model with position error:

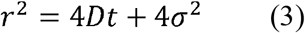

The numbers of trajectories and nuclei analyzed were: wild-type emerin (71004 trajectories, 14 nuclei); lamin A/C depletion (lamin A/C KD, 60569 trajectories,11 nuclei); nuclear actin depletion (IPO9 KD, 74501 trajectories, 17 nuclei); replacement of endogenous BAF with BAF^L58R^ (BAF KD + BAF^L58R^, 62714 trajectories, 8 nuclei); complemented wild-type emerin-GFP-emerin species assessed by CALM (4833 trajectories, 13 nuclei); wild-type emerin after nuclear deformation on 15 µm (27266 trajectories, 10 nuclei); wild-type emerin after nuclear deformation on 10 µm wide micropatterns (12915 trajectories, 8 nuclei); Q133H emerin (105050 trajectories, 13 nuclei); Δ95-99 emerin (76944 trajectories,14 nuclei); P183H emerin (86529 trajectories, 21 nuclei); and complemented P183H emerin-GFP-emerin species assessed by CALM (10519 trajectories, 21 nuclei). All diffusion coefficients *D* are reported in µm^2^ s^−1^ ± standard error of fit value (±s.e.m.). Statistical comparisons between *D* values were done using t-tests. Population percentages are derived from the averaged *a_i_(t)* values over the considered time lags.

To estimate the diffusion that might be expected for immobile PA-TagRFP-emerin (D_immobile_) under our imaging and analytical conditions, we simulated immobilized PA-TagRFP emerin with photophysical parameters similar to that of live cell acquisitions, using the software Fluosim(Lagardère et al., 2020). After PDSD analysis with SLIMfast, D_immobile_ was determined at 1.7×10^−4^ ± 2×10^−5^ μm^2^s^−1^.

Individual diffusion coefficients (*D*_i_) were obtained by fitting the individual mean square displacement (MSD) for each detected emerin over the first three time lags (*t*_1_-*t*_3_), using again a 2D Brownian diffusion model. Based on their individual *D*_i_ value, emerin trajectories were grouped into four diffusion ranges (*D*_i1_> 0.1 µm^2^ s^−1^, 0.1<*D*_i2_> 0.01 µm^2^ s^−1^, 0.1<*D*_i3_> 0.001 µm^2^ s^−1^, and *D*_i4_< 0.001 µm^2^ s^−1^) and plotted as maps.

### Spatial distribution and cluster analyses from super-resolution images

After 3D-dSTORM super-resolution imaging, the localization of individual emerin molecules and z-position assignments were performed by Gaussian fitting using rapidSTORM (version 3.3.1)(Wolter et al., 2012). Sample drift and overcounting corrections for molecules appearing in consecutive frames were done using PALMsiever(Pengo et al., 2015) and renderings of super-resolved images were done using ImageJ. Localization precisions (σ) in the x, y, and z dimensions were evaluated as previously described(Fernandez et al., 2017) and were σ_x_:8.3 nm, σ_y_:13.0 nm, and σ_z_:28.4 nm. 2D spatial pattern analyses of emerin distributions were performed on 2 µm x 2 µm regions of interest (ROI) typically chosen in NE areas having homogenous z ranges and away from the edges of nuclei, in order to limit 3D effects.

Emerin clustering was determined using an edge-corrected neighborhood density function (NDF) as previously described(Fernandez et al., 2017). Briefly, the NDF is a pairwise-correlation function similar to O-ring statistics that tallies the density of detected emerin within a ring of outer radius *r* and width *Δr* located at a distance *r* from an emerin position in the ROI and for all *r* + *Δr* in the ROI. The density of emerin as a function of distance from an average emerin was obtained with:

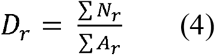

where *N_r_* is the number of neighbors and *A_r_* is the area summed over all detected emerin. NDF analyses were done over a 1µm distance on selected ROIs and with a fixed ring width of 10 nm and a ring radius increasing by 10 nm steps. To average NDF statistics from multiple ROIs across different nuclei and make them sample-size independent, *D_r_* was further standardized by dividing it by the mean density of detected emerin across the entire ROI. As such, an NDF value at a given radius indicates the relative clustering of emerin as compared to the average density across the entire sample. This relative NDF gives a value of 1 for a completely random spatial distribution as determined by Monte Carlo simulations of random emerin distributions with area and number of randomly seeded emerin equal to that of each experimental ROIs.

Relative NDF curves averaged across multiple ROIs and multiple nuclei were fitted with a previously described model(Fernandez et al., 2017), which accounts for a distribution of cluster lengths that includes two populations of emerin (monomer and oligomers) and for a probability density of emerin in 2D clusters that decays approximately as an exponential function(Sengupta et al., 2011):

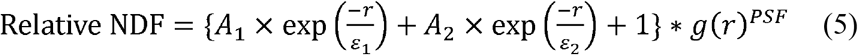

where, *A* is the clustering density, *ε* is the typical half-maximum cluster length, * denotes a 2D convolution, and *g*(*r*)*^PSF^* is the correlation function of the effective point spread function of uncertainty in position determination for the dSTORM experiments. As described previously(Fernandez et al., 2017), *g*(*r*)*^PSF^* corrects the NDF for contribution of multiple single molecule appearances (blinking) to the overall spatial distribution. After fitting relative NDF curves, the molecular density above random for emerin clusters are reported as *A* ± standard error of the fit (±s.e.m) and their typical size as 2x *ε* ± standard error of the fit and localization precision (±s.e.m). Relative increases in emerin oligomer formation during nuclear stress were determined by considering a circular shape of oligomer nanodomains and multiplying the area of oligomerization by the measured molecular density.

The numbers of localization and nuclei analyzed were: wild-type emerin in untreated cells (189331 localizations, 10 nuclei); after control siRNA transfection (n= 180546 localizations, 5 nuclei); after lamin A/C knockdown (178206 localizations, 6 nuclei); after lamin A/C knockdown and exogenous expression of lamin A/C (118859 localizations, 6 nuclei); after IPO9 knockdown (225394 localizations, 9 nuclei); after replacement of endogenous BAF by BAF^L58R^ (90,241 localizations, 6 nuclei); after SUN1 knockdown (258300 localizations, 6 nuclei); after SUN1 knockdown and exogenous expression of EGFP-SUN1 (85210 localizations, 5 nuclei); after exogenous expression of mCherry-DN-KASH (392365 localizations, 5 nuclei); after nuclear deformation on 15 µm wide micropatterns (151647 localizations, 10 nuclei); after nuclear deformation on 10 µm wide micropatterns (56563 localizations, 6 nuclei); for Q133H emerin (149340 localizations, 6 nuclei); for Δ95-99 emerin (208092 localizations, 8 nuclei); for P183H emerin (138075 localizations, 6 nuclei); for Δ95-99 emerin after nuclear deformation on 15 μm wide micropatterns (138119 localizations, 5 nuclei); and for Δ95-99 emerin after nuclear deformation on 10 μm wide micropatterns (135143 localizations, 6 nuclei).

### Cluster maps

Cluster maps were generated from drift- and overcounting-corrected super-resolved emerin positions by determining local cluster values around each emerin using the Getis and Franklin L function(Getis and Franklin, 1987) in spPack(Perry, 2004) and for a distance of 25 nm. Spatial positions in x and y, and cluster values were plotted as maps in MATLAB (MathWorks) using the meshgrid and griddata functions, a 1 nm x 1 nm pixel size and the ‘v4’ option when calculating pixel density values. The contour maps were generated using the countourf function with 200 levels. In contour maps, values *L*(r_25_)=25 represent areas where emerin is randomly distributed and *L*(r_25_)=70 values represent areas with emerin local density (70/25)^2^ = ∼8-fold higher than expected for a random distribution.

## Acknowledgements

This work was supported by the National Institute of Arthritis and Musculoskeletal and Skin Diseases of the National Institutes of Health under award number R21AR076514. We are grateful to H. Worman for providing emerin null and normal HDF and to J. Ellis, R. Frock and C. Doucet for providing emerin, lamin A/C and EGFP-SUN1 cDNAs, respectively.

