## Supplementary Information for "Emerin self-assembly and nucleoskeletal coupling regulate nuclear envelope mechanics against stress"

### SUPPLEMENTARY FIGURES

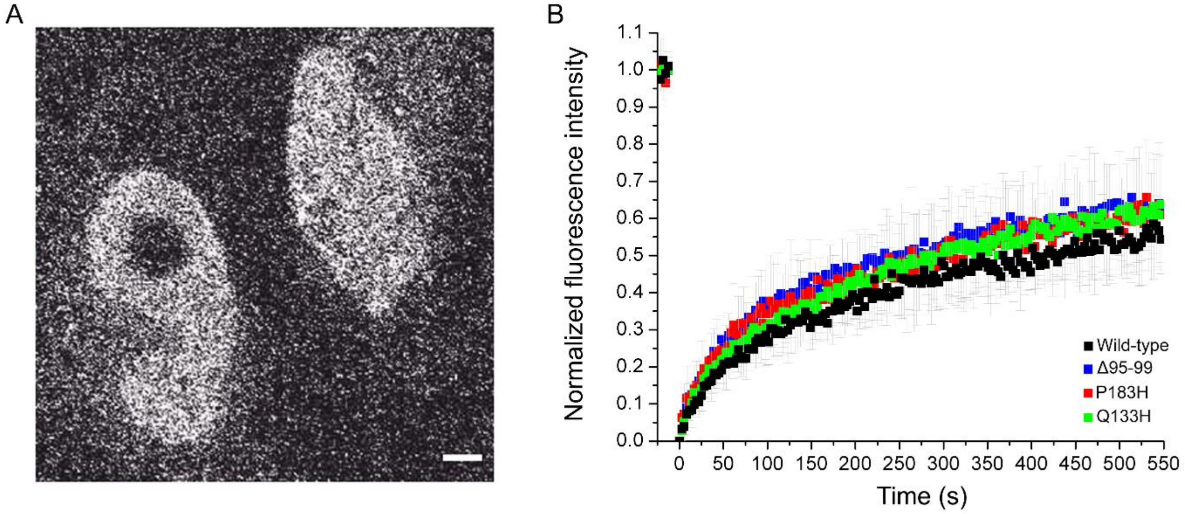


C

| | $D_{fast}$ (μm$^2$/s) | $D_{fast}$ (%) | $D_{slow}$ (μm$^2$/s) | $D_{slow}$ (%) |
|---|---|---|---|---|
| **Wildtype** | $4.48 \times 10^{-2} \pm 1 \times 10^{-2}$ | 17 | $4.0 \times 10^{-3} \pm 4 \times 10^{-4}$ | 83 |
| **P183H** | $7.73 \times 10^{-2} \pm 2 \times 10^{-2}$ | 17 | $5.0 \times 10^{-3} \pm 5 \times 10^{-4}$ | 83 |
| **Q133H** | $9.97 \times 10^{-2} \pm 3 \times 10^{-2}$ | 9 | $6.7 \times 10^{-3} \pm 3 \times 10^{-4}$ | 91 |
| **Δ95-99** | $5.22 \times 10^{-2} \pm 1 \times 10^{-2}$ | 32 | $4.7 \times 10^{-3} \pm 5 \times 10^{-4}$ | 68 |

**Figure S1. FRAP of emerin in U2OS cells**. (A) U2OS cells transfected with pA-TagRFP-emerin fusions after photoactivation and photobleaching of a circular region of interest at the nuclear envelope. Scale bar: 5 µm. (B) FRAP recovery curves of wild-type (WT), P183H, Q133H, and Δ95-99 mutated emerin at the nuclear envelope. Grey bars represent the standard deviation of the mean at each time points. n= 9-12 cells per condition. (C) Diffusion coefficients of wild-type and mutated emerin determined by fitting FRAP recovery curves with a two-components fit.

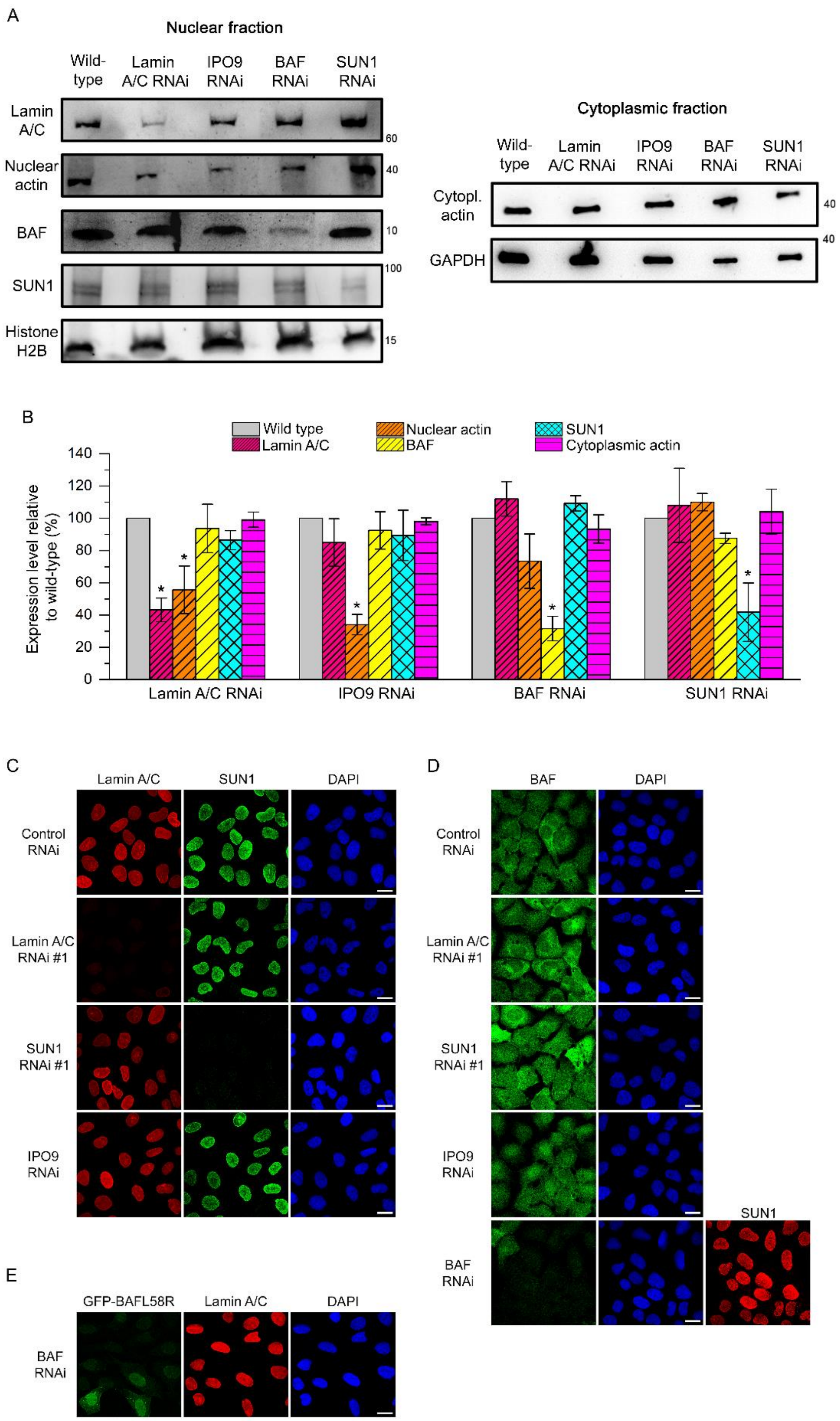


**Figure S2. Immunoblotting and immunostaining to assess the direct and indirect effects of siRNA knockdown against lamin A/C, IPO9, BAF and SUN1.** (A) Western blots of lamin A/C, nuclear actin, BAF, SUN1 and histone H2A in the nuclear fraction (left) and of actin and GAPDH in the cytoplasmic fraction (right) of *EMD*$^{+/y}$ human dermal fibroblasts after RNA interference**.** (B) Relative change in lamin A/C, nuclear actin, BAF, SUN1 and cytoplasmic actin expression levels compared to untreated cells after siRNA **(**t-test, *: $p<0.05$)**.** Error bars represent the standard deviation of the mean**.** (C) Confocal fluorescence imaging of lamin A/C, SUN1 and the nucleus (DAPI) after siRNA-induced depletion of lamin A/C, SUN1 and IPO9. All scale bars: 20 µm. (D) Confocal fluorescence imaging of endogenous BAF, the nucleus (DAPI) and SUN1 after siRNA-induced depletion of lamin A/C, SUN1, IPO9 or BAF. All scale bars: 20 µm. (E) Confocal fluorescence imaging of GFP-BAF$^{L58R}$, lamin A/C and the nucleus (DAPI) after siRNA-induced depletion of endogenous BAF. Scale bar: 20 µm.

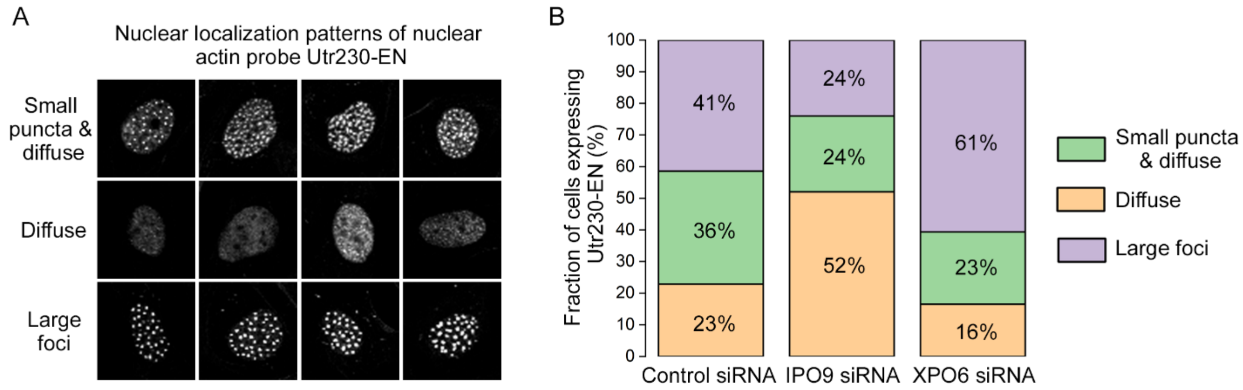


**Figure S3. Effects of IPO9 siRNA and XPO6 siRNA on nuclear actin organization**. (A) Examples of nuclear localization patterns for the short nuclear actin filament probe Utr230-EN in HDF cells. Patterns are classified as: (i) small puncta and diffuse, (ii) diffuse or (iii) large foci, reflecting variations in nuclear actin filament content across cells. (B) Distribution of nuclear actin filament classes after control siRNA treatment (n = 635 nuclei), IPO9 siRNA (n = 663 nuclei) or XPO6 siRNA (n = 625 nuclei). Knockdown of IPO9 results in the majority of cells displaying a diffused Utr230-EN pattern, indicative of lower nuclear actin filament contents. Inversely, knockdown of XPO6 results in the majority of cells displaying larger and brighter foci compared to control siRNA, indicative of increased nuclear actin filament contents.

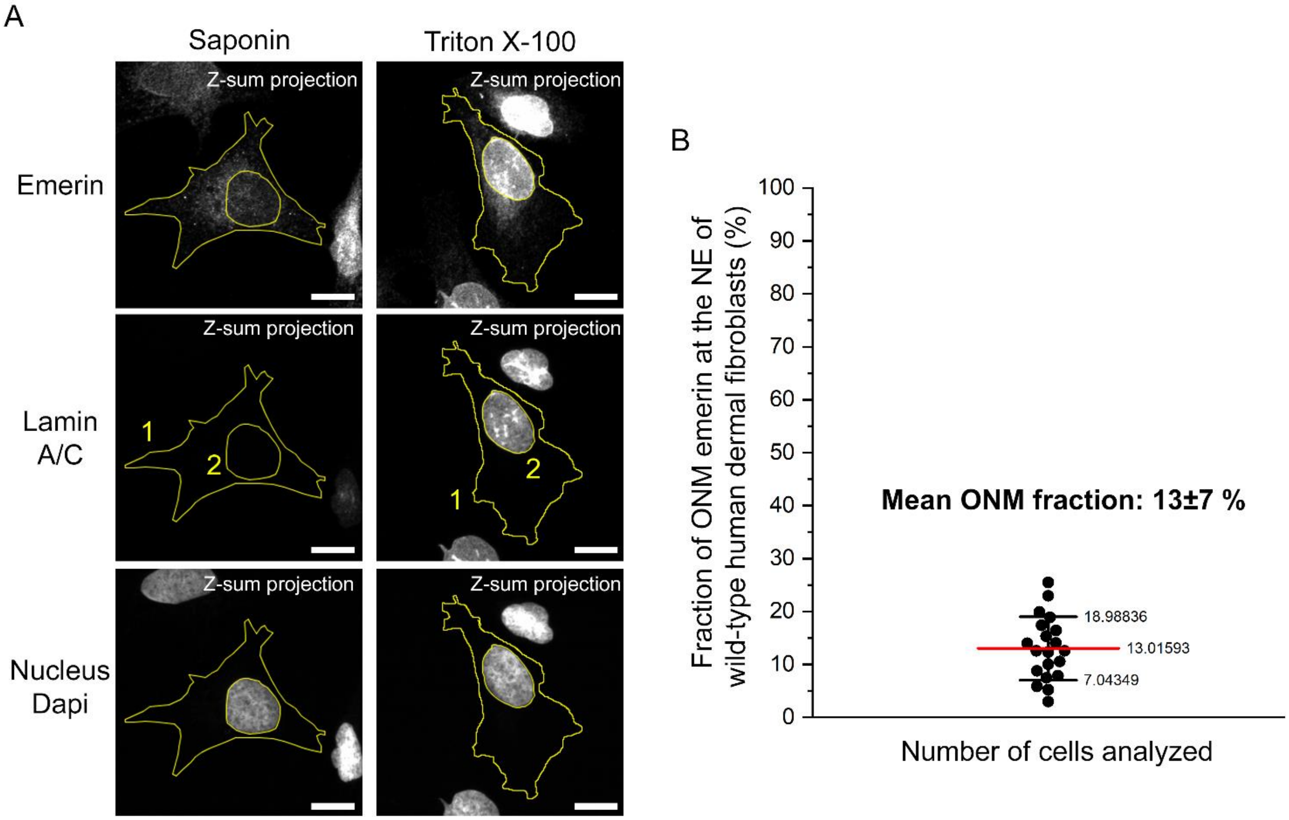


**Figure S4. Quantification of ONM emerin fraction in *EMD*$^{+/y}$ human dermal fibroblasts from confocal z-stacks**. (A) Example of z-stack sum intensity projections for fibroblasts permeabilized with saponin or triton X-100 and immunostained for emerin, lamin A/C and Dapi. Region of interests (ROIs) used to measure fluorescence intensities from emerin immunostaining are shown in yellow. Scale bars: 15 µm. (B) Histogram of emerin ONM fraction at the nuclear envelope determined from confocal z-stack projections and nuclear ROI quantifications of saponin treated cells (N=20) and Triton X-100 treated cells (N=20). Mean is indicated by the red bar.

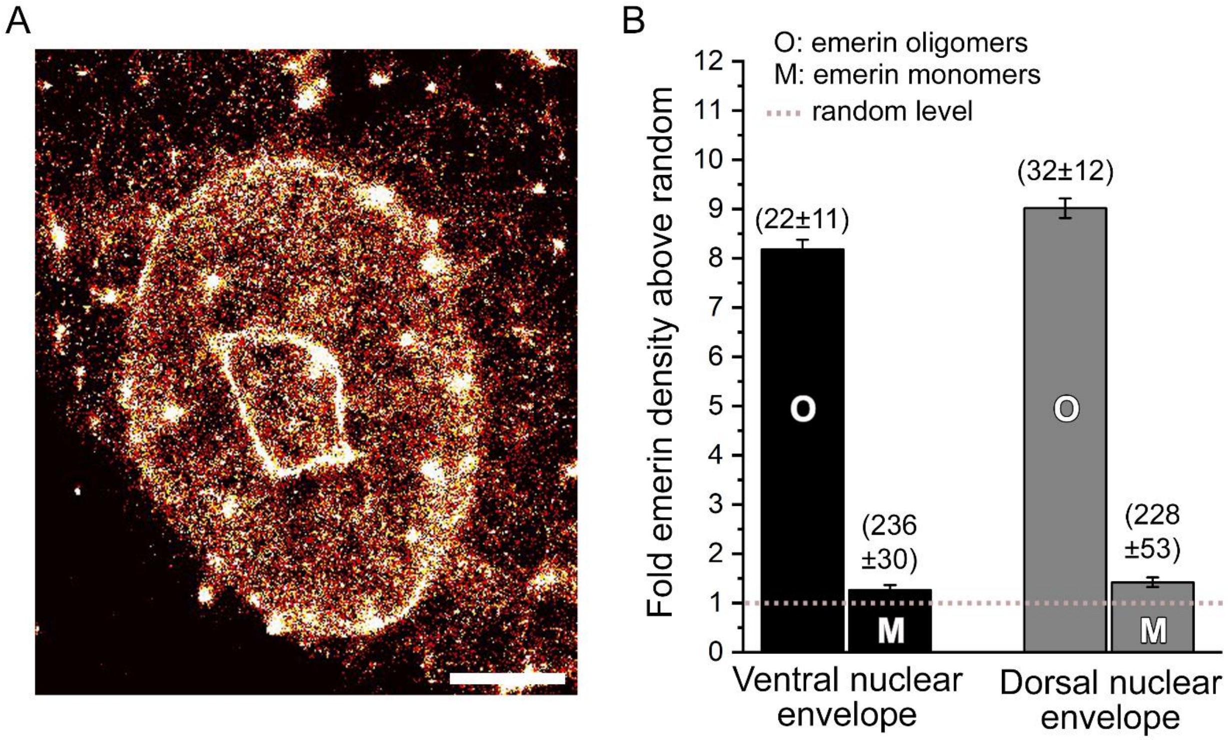


**Figure S5. Nanoscale organization of emerin at the dorsal nuclear envelope of human dermal fibroblasts**. (A) Two-dimensional rendering of wild-type SNAP-emerin imaged by 3D super-resolution at the dorsal nuclear envelope. Areas with very high densities of emerin (white spots and lines) correspond to local nuclear envelope folds often observed in the plane of the dorsal nuclear envelope and to the 2D-projection of different z-positions of emerin at the nuclear rim. Scale: 5 µm. (B) Comparison of molecular densities above random (± s.e.m.) for wild-type emerin oligomers (O) and monomers (M) at the ventral (189331 localizations, 10 nuclei) or the dorsal (51387 localizations, 4 nuclei) nuclear envelope of human dermal fibroblasts. Values in parenthesis represent the size (± s.e.m.) of each domain in nanometers.

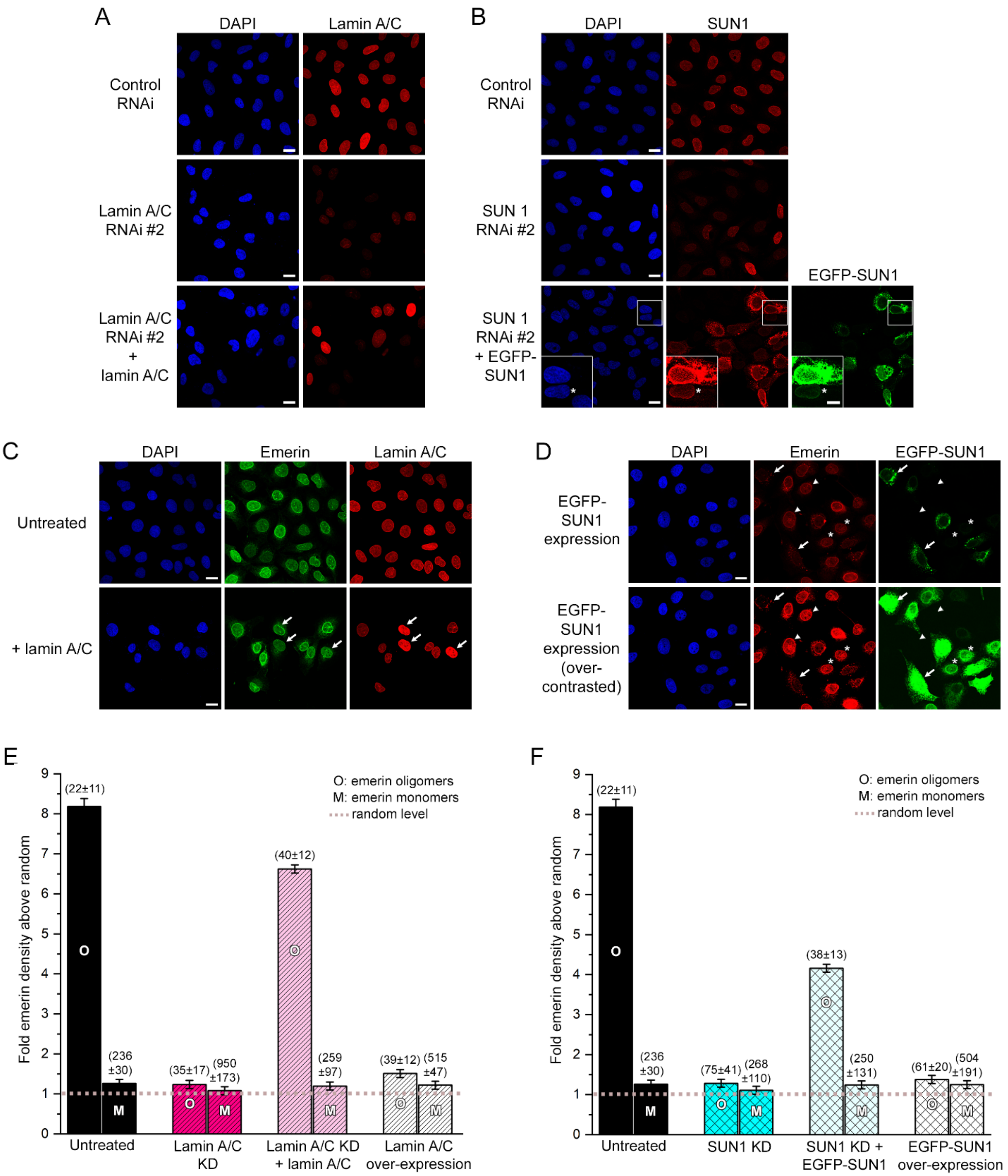


**Figure S6. Rescue by expression of exogenous lamin A/C or SUN1 after siRNA knock-down and effects of their over-expression on emerin localization and nanoscale organization**. A) Confocal fluorescence imaging of lamin A/C and the nucleus (DAPI) after siRNA-induced depletion of lamin A/C and re-expression of exogenous and siRNA-resistant lamin A/C. Scales: 20 µm. B) Confocal fluorescence imaging of SUN1, EGFP-SUN1 and the nucleus (DAPI) after siRNA-induced depletion of endogenous SUN1 and re-expression of exogenous and siRNA-resistant EGFP-SUN1. The over-contrasted inset shows that overexpression of EGFP-SUN1 induces the mis-localization of SUN1 from the nuclear envelope, while cells with low EGFP-SUN1 expression display relatively normal nuclear envelope accumulation (star). Scales: 20 µm, inset: 10 µm. C) Confocal fluorescence imaging of lamin A/C, emerin and the nucleus (DAPI) after overexpression of exogenous lamin A/C. Cells overexpressing lamin A/C (arrows) display an apparently normal emerin localization at the nuclear envelope. Scales: 20 µm. D) Confocal fluorescence imaging of emerin, EGFP-SUN1 and the nucleus (DAPI) after overexpression of exogenous EGFP-SUN1. EGFP-SUN1 overexpression induces a mis-localization of emerin from the nuclear envelope (arrows), while low EGFP-SUN1 expression levels (stars) result in relatively normal emerin localization at the nuclear envelope compared to non-expressing cells (arrowheads). Scales: 20 µm. (E) Molecular densities above random (± s.e.m.) for wild-type emerin in untreated cells, after lamin A/C knock down, lamin A/C knock down and exogenous expression of lamin A/C (118859 localizations, 6 nuclei) or overexpression of lamin A/C (204532 localizations, 5 nuclei). Values in parenthesis represent the size (± s.e.m.) of each domain in nanometers. (F) Molecular densities above random (± s.e.m.) for wild-type emerin in untreated cells, after SUN1 knock down (258300 localizations, 6 nuclei), SUN1 knock down and exogenous expression of EGFP-SUN1 (85210 localizations, 5 nuclei) or overexpression of EGFP-SUN1 (288522 localizations, 7 nuclei). Values in parenthesis represent the size (± s.e.m.) of each domain in nanometers.

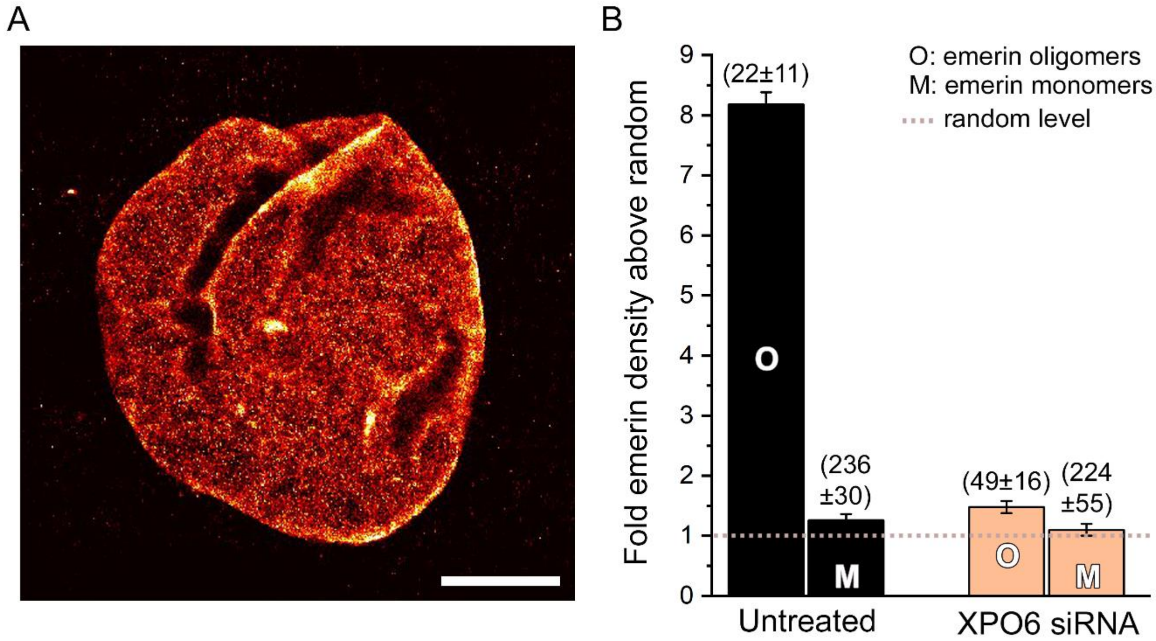


**Figure S7. Nanoscale organization of wild-type emerin following increase in nuclear actin levels by siRNA of exportin-6.** (A) Two-dimensional rendering of wild-type SNAP-emerin imaged by 3D super-resolution in HDF cells treated with siRNA against exportin-6 (XPO6). Some nuclei appeared locally crumpled after siRNA of XPO6. Scale: 5 µm. (B) Molecular densities above random (± s.e.m.) for wild-type emerin oligomers (O) and monomers (M) in untreated cells or after XPO6 knock down to increase nuclear actin levels (315527 localizations, 7 nuclei). Values in parenthesis represent the size (± s.e.m.) of each domain in nanometers.

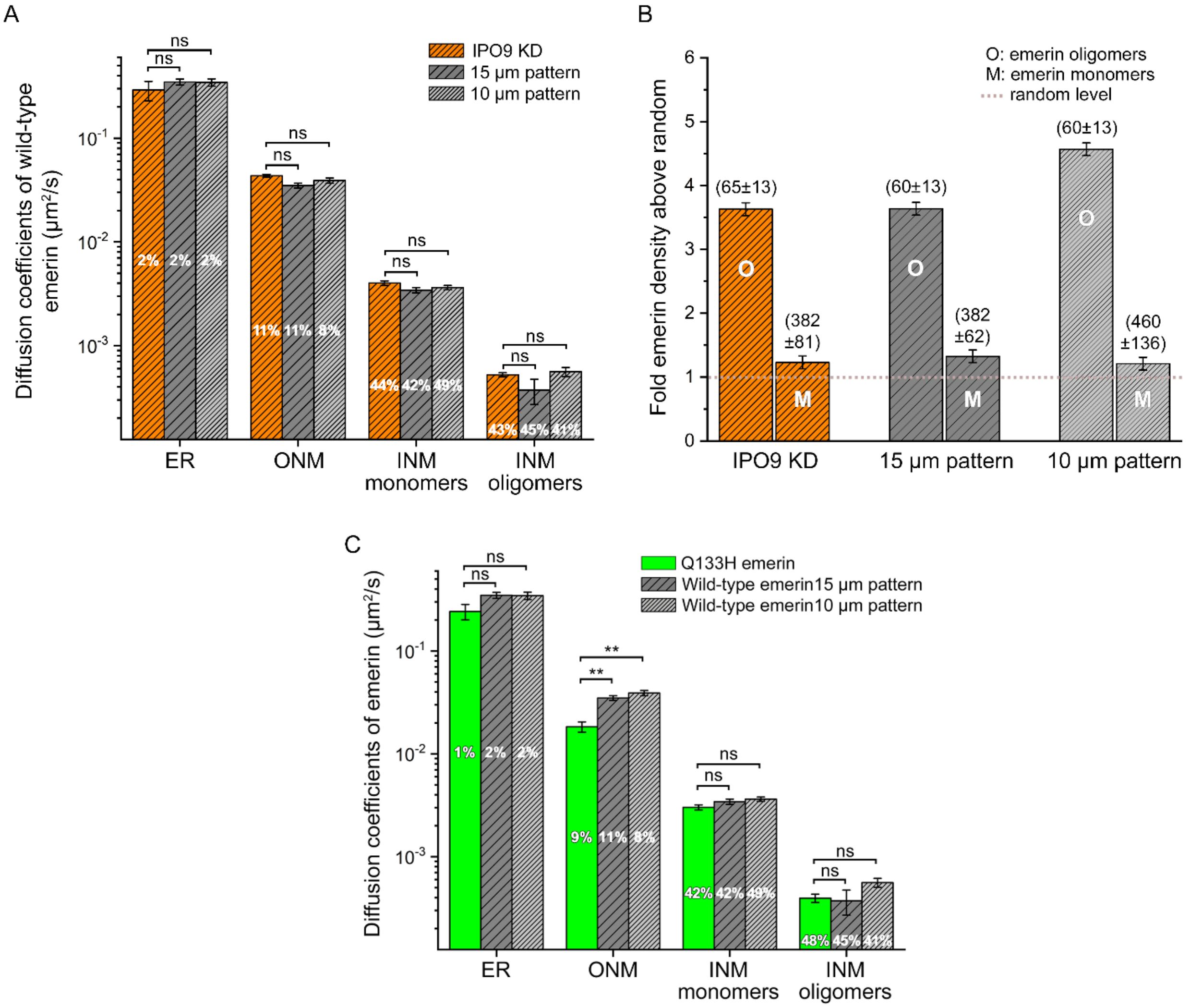


**Figure S8. Comparison of wild-type emerin organization after nuclear actin depletion or cell micropatterning and of Q133H and wild-type emerin diffusive behaviors.** (A) Diffusion coefficients (± s.e.m.) and population percentage of wild-type emerin after nuclear actin depletion by IPO9 knock down (IPO9 KD, *n*=74501trajectories in 17 nuclei), after nuclear deformation on 15 µm wide (*n n*=27266 trajectories in 10 nuclei) or after nuclear deformation 10 µm wide micropatterns (*n*=12915 trajectories in 8 nuclei, t-test, ns: non-significant). (B) Relative nuclear envelope molecular densities above random (± s.e.m.) for wild-type emerin oligomers (O) and monomers (M) in non-patterned *Emd*$^{-/y}$ fibroblasts after nuclear actin depletion (IPO9 KD, n=225,394 localizations in 9 nuclei) and in micropatterned fibroblasts with deformed nucleus on 15 µm wide (n=151,647 localizations in 10 nuclei) or 10 µm wide micropatterns (n=56,563 localizations in 6 nuclei). Values in parenthesis represent the typical length size (± s.e.m.) of each domain in nanometers. (C) Comparison of Q133H emerin diffusion (± s.e.m.) and population percentage in non-patterned cells with that of wild-type emerin under mechanical stress on micropatterns (t-test, ns: non-significant, **: $p$<0.01).

### SUPPLEMENTARY TABLES

**Table S1.** Diffusion coefficients of wild-type and mutated emerin determined by PDSD analyses after sptPALM or CALM imaging.

| Emerin | $D_1$ (μm²/s) | $D_2$ (μm²/s) | $D_3$ (μm²/s) | $D_4$ (μm²/s) |
|---|---|---|---|---|
| **Wild-type** | $2.21x10^{-1}±4.9x10^{-2}$ | $1.48x10^{-2}±1.5x10^{-3}$ | $1.73x10^{-3}±1.1x10^{-4}$ | $2.6x10^{-4}±1x10^{-5}$ |
| **Wild-type split-GFP complemented** | not detected | $1.93x10^{-2}±2.5x10^{-3}$ | not detected | $3.1x10^{-4}±5x10^{-5}$ |
| **Wild-type + lamin A/C knock down** | $2.24x10^{-1}±6.1x10^{-2}$ | $1.94x10^{-2}±1.9x10^{-3}$ | $3.94x10^{-3}±4.1x10^{-4}$ | $4.5x10^{-4}±3x10^{-5}$ |
| **Wild-type + IPO9 knock down** | $2.91x10^{-1}±6.9x10^{-2}$ | $4.34x10^{-2}±2.7x10^{-3}$ | $4.01x10^{-3}±2.0x10^{-4}$ | $5.2x10^{-4}±2x10^{-5}$ |
| **Wild-type + BAF L58R** | $3.13x10^{-1}±5.3x10^{-2}$ | $5.60x10^{-2}±1.4x10^{-3}$ | $9.29x10^{-3}±9.3x10^{-4}$ | $9.6x10^{-4}±6x10^{-5}$ |
| **Wild-type in 15 μm pattern** | $3.48x10^{-1}±2.3x10^{-2}$ | $3.49x10^{-2}±1.8x10^{-3}$ | $3.43x10^{-3}±2.0x10^{-4}$ | $3.7x10^{-4}±9x10^{-5}$ |
| **Wild-type in 10 μm pattern** | $3.45x10^{-1}±2.8x10^{-2}$ | $3.90x10^{-2}±2.3x10^{-3}$ | $3.63x10^{-3}±1.8x10^{-4}$ | $5.6x10^{-4}±5x10^{-5}$ |
| **Q133H** | $2.42x10^{-1}±4.1x10^{-2}$ | $1.83x10^{-2}±2.1x10^{-3}$ | $3.02x10^{-3}±1.7x10^{-4}$ | $3.9x10^{-4}±4x10^{-5}$ |
| **Δ95-99** | $2.58x10^{-1}±1.1x10^{-2}$ | $1.41x10^{-2}±1.3x10^{-3}$ | $1.16x10^{-3}±0.6x10^{-4}$ | $1.6x10^{-4}±1x10^{-5}$ |
| **P183H** | $2.16x10^{-1}±2.1x10^{-2}$ | $1.34x10^{-2}±1.8x10^{-3}$ | $1.13x10^{-3}±1.3x10^{-4}$ | $0.9x10^{-4}±2x10^{-5}$ |
| **P183H split-GFP complemented** | $0.97x10^{-1}±1.6x10^{-2}$ | $1.15x10^{-2}±3.0x10^{-3}$ | $1.37x10^{-3}±1.8x10^{-4}$ | not detected |

**Table S2.** Molecular densities and domain sizes of wild-type and mutated emerin determined by spatial distribution analyses of emerin in super-resolved images.

| Emerin | Oligomer density (fold above random) | Oligomer domain size (nm) | Monomer density (fold above random) | Monomer domain size (nm) |
|---|---|---|---|---|
| **Wild-type** | 8.2±0.2 | 22±11 | 1.3±0.1 | 236±30 |
| **Wild-type + siRNA control** | 7.4±0.2 | 19±12 | 1.3±0.1 | 205±53 |
| **Wild-type + lamin A/C knock down** | 1.2±0.1 | 35±17 | 1.1±0.1 | 950±173 |
| **Wild-type + lamin A/C knock down + lamin A/C** | 6.6±0.1 | 40±12 | 1.2±0.1 | 259±97 |
| **Wild-type + lamin A/C overexpression** | 1.5±0.1 | 39±12 | 1.2±0.1 | 515±47 |
| **Wild-type + IPO9 knock down** | 3.6±0.1 | 65±13 | 1.2±0.1 | 382±81 |
| **Wild-type + XPO6 knock down** | 1.5±0.1 | 49±16 | 1.2±0.1 | 224±55 |
| **Wild-type + BAF L58R** | 2.0±0.1 | 85±19 | 1.3±0.1 | 718±250 |
| **Wild-type + SUN 1 knock down** | 1.2±0.1 | 75±41 | 1.1±0.1 | 268±110 |
| **Wild-type + SUN 1 knock down + EGFP-SUN1** | 4.2±0.1 | 38±13 | 1.2±0.1 | 250±131 |
| **Wild-type + EGFP-SUN1 over-expression** | 1.4±0.1 | 61±20 | 1.2±0.1 | 504±191 |
| **Wild-type + DN-KASH domain** | 1.2±0.1 | 61±20 | 1.1±0.1 | 393±119 |
| **Wild-type in 15 µm pattern** | 3.6±0.1 | 60±13 | 1.3±0.1 | 382±62 |
| **Wild-type in 10 µm pattern** | 4.6±0.1 | 60±13 | 1.2±0.1 | 460±136 |
| **Q133H** | 12.2±0.2 | 19±12 | 1.3±0.1 | 213±62 |
| **Δ95-99** | 1.3±0.1 | 48±14 | 1.1±0.1 | 420±51 |
| **P183H** | 2.2±0.1 | 35±12 | 1.3±0.1 | 321±29 |
| **Δ95-99 in 15 µm pattern** | 1.7±0.1 | 81±16 | 1.2±0.1 | 810±215 |
| **Δ95-99 in 10 µm pattern** | 2.0±0.1 | 75±20 | 1.2±0.1 | 499±250 |

### SUPPLEMENTARY MOVIES

**Movie S1.** sptPALM imaging of wild-type PA-TagRFP-emerin at the nuclear envelope of an emerin-null dermal fibroblast (*Emd*$^{-/y}$). Point spread function have been intentionally expended to facilitate visualization. Movie acquired at 40 ms/frame and played back at the same rate.

**Movie S2.** CALM imaging of complemented wild-type sGFP-emerin and M3-emerin dimers and oligomers at the nuclear envelope of an emerin-null dermal fibroblast (*Emd*$^{-/y}$). Movie acquired at 40 ms/frame and played back at the same rate.

**Movie S3.** sptPALM imaging of wild-type PA-TagRFP-emerin at the nuclear envelope of an emerin-null dermal fibroblast (*Emd*$^{-/y}$) micropatterned on a 15 µm wide fibronectin strip. Movie acquired at 40 ms/frame and played back at the same rate.
